# Identification and characterisation of *Klebsiella pneumoniae* and *Pseudomonas aeruginosa* clinical isolates with atypical β-lactam susceptibility profiles using Orbitrap liquid chromatography-tandem mass spectrometry

**DOI:** 10.1101/2022.02.27.482154

**Authors:** Yuiko Takebayashi, Punyawee Dulyayangkul, Naphat Satapoomin, Wan Ahmad Kamil Wan Nur Ismah, O. Martin Williams, Alasdair P. Macgowan, Kate J. Heesom, Philip B. Williams, Matthew B. Avison

## Abstract

There is significant interest in the possibility of predicting antibacterial drug susceptibility directly though the analysis of bacterial DNA or protein. We report the use of *Klebsiella pneumoniae, Escherichia coli, Pseudomonas aeruginosa* and *Acinetobacter baumannii* transformants to define baseline predictive rules for the β- lactam susceptibility profiles of β-lactamase positive clinical isolates. We then deployed a robust and reproducible shotgun proteomics methodology to identify β-lactamase positivity and predict β-lactam susceptibility by reference to our baseline predictive rules both in cultured bacteria and in extracts of culture-positive blood. Proteomics and whole genome sequencing then allowed us to characterise *K. pneumoniae* and *P. aeruginosa* isolates that differed from the expected β-lactam susceptibility profile, iteratively expanding our predictive rules. Proteomics added considerable value over and above the information generated by whole genome sequencing, allowing for gene expression, not just gene presence to be considered. Specifically, in *K. pneumoniae*, we identified key differences between *acrR* and *ramR* regulatory mutations and compared the effects of OmpK36 Aspartate-Threonine or Glycine-Aspartate dipeptide porin insertions on susceptibility to cefepime and carbapenems. In *P. aeruginosa*, we identified differences in the gene expression effects of *mexR* versus *nalC* mutations and related these to differences in β-lactam MICs against isolates hyper-producing AmpC β-lactamase and or producing a metallo-β-lactamase.

## Introduction

One way to dramatically reduce the time it takes to provide key information relevant to antibiotic choice is to identify antibiotic resistance (ABR) mechanisms from clinical samples. PCR can be used to identify certain mobile ABR genes. A wider variety of ABR genes might be identified in one assay using microarray hybridisation technology, but whole genome sequencing (WGS) is touted as being a more comprehensive answer to this question (1-6). In recent years, the speed, accuracy, cost, and the amount of DNA required for WGS have all shifted by orders of magnitude in favour of routine WGS from clinical samples and some major successes have been recorded, particularly where bacterial density is high, e.g., urinary tract infection (7). Importantly, WGS potentially allows the complex interplays between mobile ABR genes and background mutations to be integrated in a prediction of ABR, something that is particularly necessary in Gram-negative bacteria, where ABR phenotype is frequently multi-factorial (8). However, this highlights our lack of understanding of the way genotype relates to ABR phenotype. Without a detailed understanding of which mutations influence, and which do not influence ABR, we cannot hope to accurately predict ABR from WGS in all cases. The need for more research in this area was highlighted in a recent EUCAST sub-committee report (9).

One of the main information weaknesses of using WGS is a lack of understanding about how genotype affects gene expression levels; for ABR, protein abundance can have a profound effect. For example, in the presence of weak carbapenem hydrolysing β-lactamases, such as CTX-M and CMY, mutations that reduce the rate of carbapenem entry into Enterobacterales isolates can help confer carbapenem resistance (10-13).

In an attempt to mitigate this weakness, we have established a “shotgun” proteomics methodology using a nano-liquid chromatography, Orbitrap tandem mass spectrometry (LC-MS/MS) approach. This has been used to characterise proteomic responses to mutations in regulators that affect ABR (13). Based on the impacts of these mutations on protein production, this approach has also been used to identify novel mutant alleles in ABR regulators in clinical isolates grown in culture, thereby enhancing the power of WGS to predict fluoroquinolone susceptibility (14).

In the work reported here, we have used our LC-MS/MS methodology to identify and quantify acquired β-lactamases and known, intrinsic ABR proteins directly from bacterial isolates grown in culture and directly from culture positive blood. We have used our analysis of wild-type backgrounds to establish baseline rules for the prediction of β-lactam susceptibility across a range of Gram-negative species. We then show how the method can be used to identify clinical isolates that differ from this baseline prediction, allowing those “atypical” isolates to be targeted for experimentation to learn more about the biology of ABR.

## Results

### Establishing baseline rules for predicting β-lactam resistance in Gram negative bacteria

The predominant mechanism of clinically important β-lactam resistance in Gram-negative bacteria is the production of β-lactamase enzymes (12). Our first aim was to define baseline rules that would allow us to predict the β-lactam resistance profiles of bacteria producing different β-lactamases. To do this, seven of the most clinically important β-lactamase genes were cloned downstream of a natural promoter (representative of promoters found in clinical isolates) and expressed in four clinically relevant test species (*Klebsiella pneumoniae, Escherichia coli, Pseudomonas aeruginosa* and *Acinetobacter baumannii*). The β-lactam susceptibility profile of each transformant was then determined (**Table S1**).

We therefore established crude baseline predictive rules: if species X produces β-lactamase Y it will have antibiogram defined in **Table S1**. Real-world bacterium to bacterium variation in β-lactamase abundance and the abundance of proteins that influence background β-lactam accumulation are likely to affect the general applicability of these baseline predictive rules.

Our ultimate objective was to factor in such variation by using LC-MS/MS proteomics. But first, we needed to confirm that our proteomics method can accurately and reproducibly quantify proteins relevant to β-lactam resistance, and we did this in our seven β-lactamase gene transformants. Our methodology allows identification and quantification of >1000 proteins in each sample tested. **Table 1** reports abundance values (normalised to the average abundance of ribosomal proteins) for specific, relevant ABR proteins from each of the *E. coli* transformants grown in liquid culture. They were easily identified when running 20 μg of total protein and the relative abundance of each chromosomally encoded porin or efflux pump protein was very similar from run to run (*n*=3 biological replicates for each transformant). This confirmed the reproducibility of our method. To test the limit of detection, total protein derived from four of the *E. coli* transformants (KPC-3, VIM-1, CMY-2, CTX-M-15) was mixed, with each in approximately equal amounts and diluted in 10-fold serial dilution. Samples were then analysed using 4 μg, 400 ng or 40 ng of total mixed protein. All ABR proteins were successfully identified, even when 40 ng of total protein was analysed from a complex mixture. The observed absolute abundance for each protein in the mixture followed the expected 10-fold serial reduction, confirming the accuracy of our method (**Table 2**).

**Table 1.**
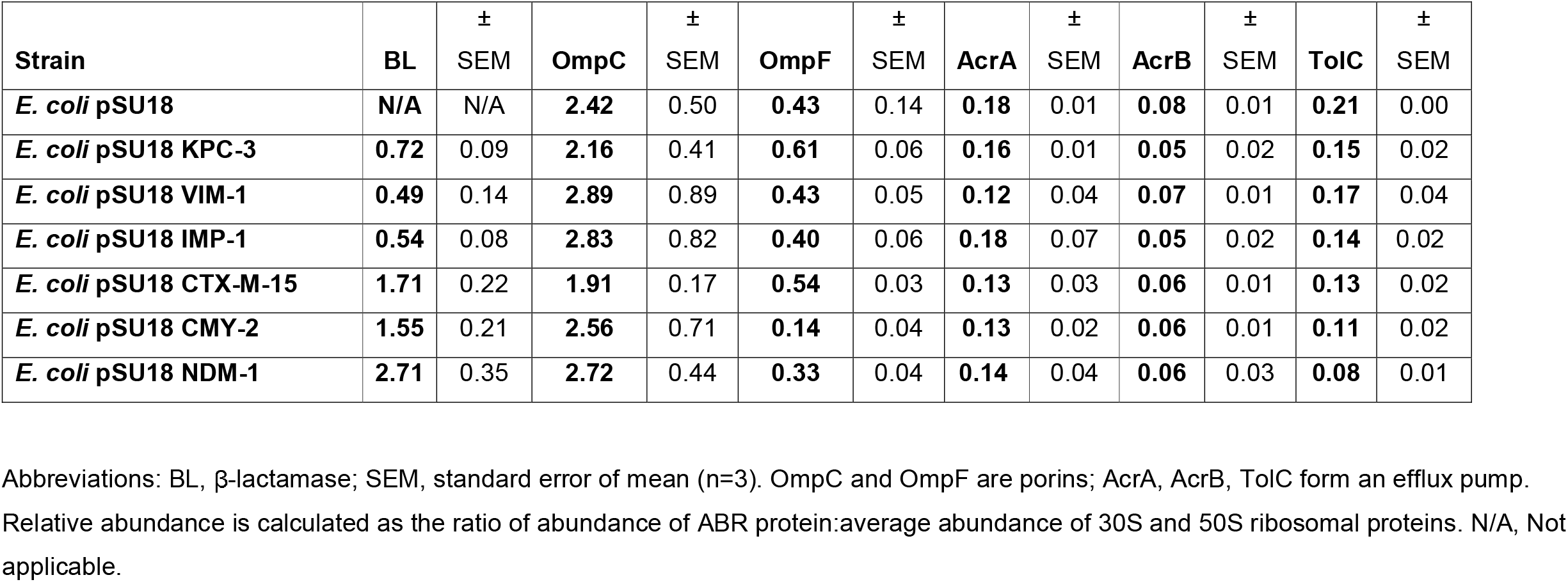
Relative abundance of ABR proteins in *E. coli* β-lactamase transformants.

**Table 2.** Abundance of ABR proteins in mixtures of protein from four *E. coli* transformants measured using various total protein amounts.

| **Amount of Total Protein in Mixture** | **KPC-3** | **VIM-1** | **CMY-2** | **CTX-M-15** | **OmpA** | **OmpC** | **OmpF** | **AcrA** | **AcrB** | **TolC** | **Average RP** |
| --- | --- | --- | --- | --- | --- | --- | --- | --- | --- | --- | --- |
| **4 µg** | 10.50 | 10.63 | 34.49 | 18.50 | 167.55 | 236.78 | 58.73 | 16.41 | 4.55 | 11.95 | **97.49** |
| **0.4 µg** | 0.87 | 0.85 | 2.79 | 1.45 | 19.43 | 21.18 | 5.63 | 0.92 | 0.50 | 0.84 | **18.03** |
| **0.04 µg** | 0.08 | 0.05 | 0.14 | 0.07 | 1.82 | 1.68 | 0.56 | 0.07 | 0.01 | 0.07 | **0.72** |

Abbreviations: RP = ribosomal protein. Abundance values represented as quotients of $10^7$ arbitrary units. n=1 for each run.

### Identifying K. pneumoniae clinical isolates that differ from the β-lactam resistance pattern predicted using our baseline predictive rules

Because we did not understand the relationships between β-lactamase, efflux pump and porin abundance (as defined using our LC-MS/MS methodology) and β-lactam susceptibility, our initial baseline predictive rules to define β-lactam susceptibility for this work were binary: if a β-lactamase was identified in an isolate of a particular species, then susceptibility was predicted simply by reference to our baseline predictive rules (**Table S1**). These rules were first used to predict cephalosporin and carbapenem susceptibility in 40 *K. pneumoniae* clinical isolates grown in broth culture as shown in **Table 3**.

**Table 3.**
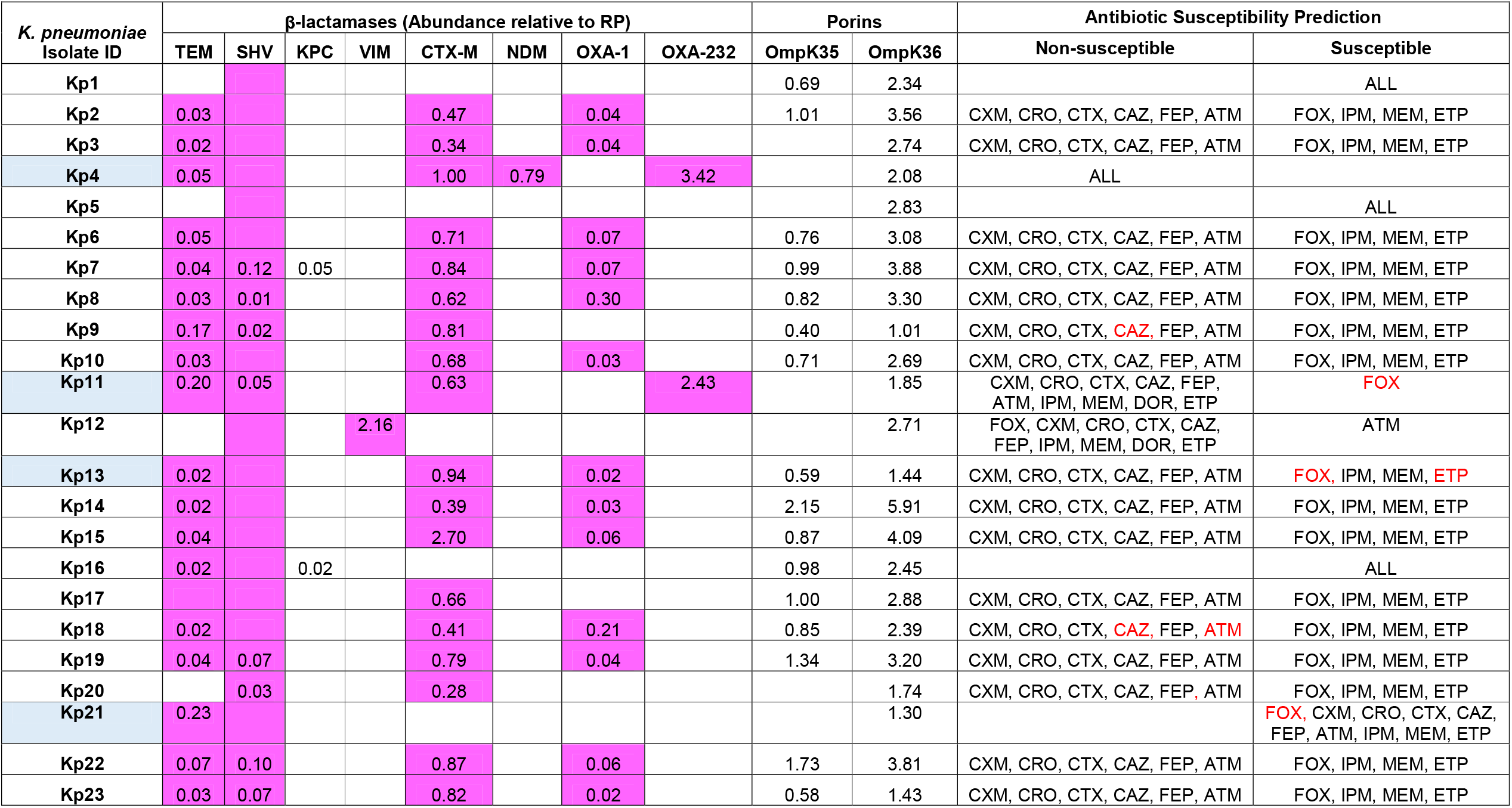
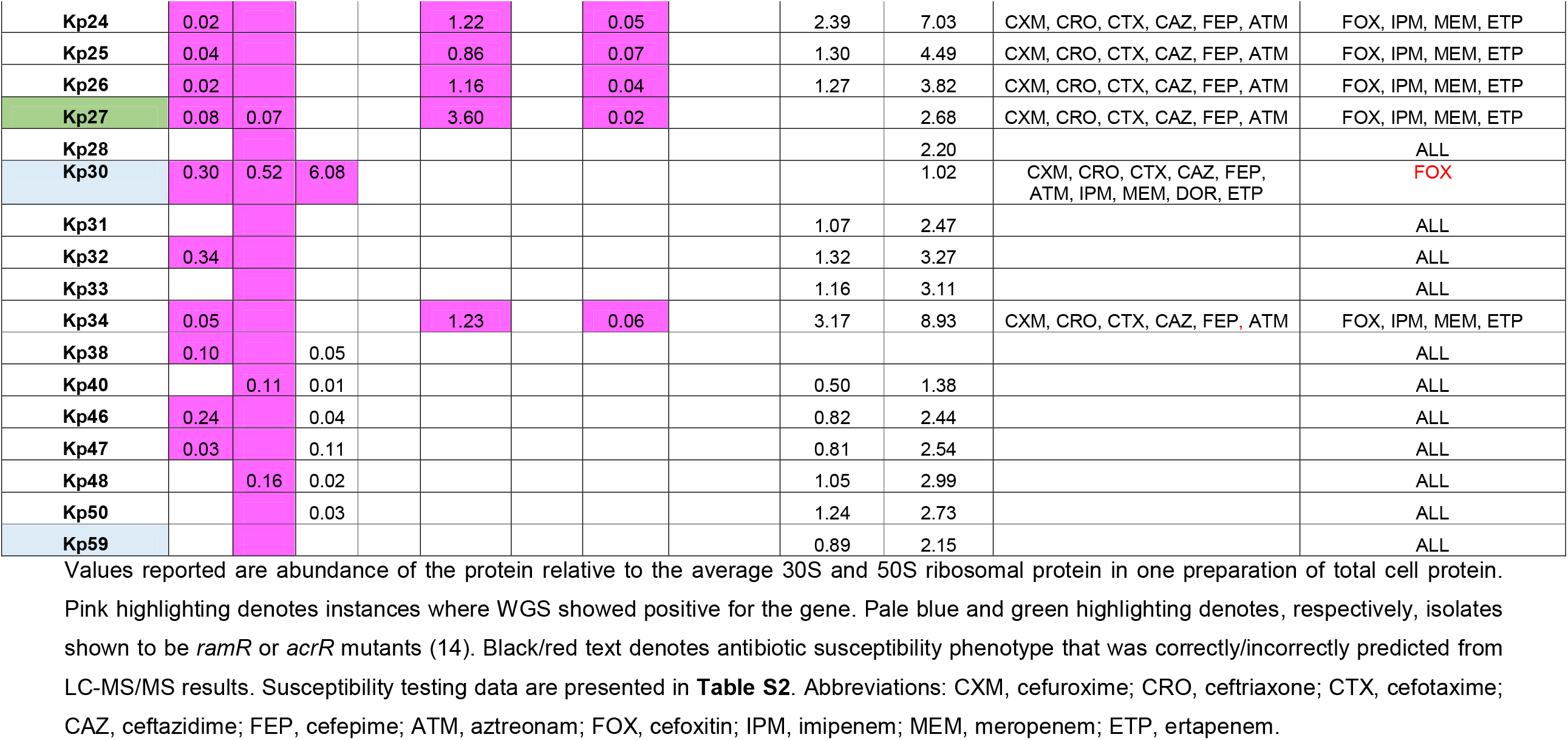
Prediction of β-lactam susceptibility in *K. pneumoniae* clinical isolates

WGS agreed with the binary calling of β-lactamase complement by LC-MS/MS in all 40 isolates with two exceptions (**Table 3**). SHV β-lactamase is intrinsic to *K. pneumoniae* (18) and all isolates were positive based on WGS, as expected, but few were positive in the LC-MS/MS data. It is well known that basal transcription from the chromosomal *bla*_SHV_ promoter is usually very low (15) and so it was not surprising that protein levels were below the limit of detection in many cases. In 25% of isolates, KPC β-lactamase was detected though only in one case was this confirmed by WGS (**Table 3**). This disparity likely arises because there is some protein/s in these samples having a few peptides with significant identity to peptides from KPC. It is important to note that ours is a shotgun LC-MS/MS approach, designed to be used in conjunction with WGS, and so it can identify and quantify hundreds of proteins. By more judiciously selecting target peptides representative of ABR proteins it is certainly possible to increase specificity, as has been shown previously (16). Indeed, the only true KPC positive sample, as validated by WGS, matched to 26 peptides of KPC, where the false positives matched to fewer than 6 peptides. So, this provides a simple rationale for simply excluding false positives for KPC production.

Analysis of these proteomics and WGS data in conjunction with susceptibility testing assays (**Table S2**) confirmed that β-lactam susceptibility was correctly predicted in 34/40 of the *K. pneumoniae* isolates (**Table 3**) based on our baseline predictive rules (**Table S1**).

### Identification of phenotypic differences associated with ramR and acrR mutation in K. pneumoniae

Critical errors, where we predicted β-lactam susceptibility in *K. pneumoniae*, but the isolate was non-susceptible, were seen for at least one β-lactam in 4/40 isolates – KP11, KP13, KP21 and KP30 – suggesting that these isolates are unusual, and worthy of additional study (**Table 3**). Detailed analysis of the proteomics data revealed that the distribution curve of average AcrA/AcrB efflux pump abundance for the 40 *K. pnuemoniae* isolates was clear and all four of the isolates with critical errors fall to the right of the “wild-type” distribution (**Fig. S1**). AcrAB-TolC over-production has previously been shown to enhance β-lactam resistance (13) so we suspected that this might explain the critical errors in these four isolates. However, three other isolates, KP4, KP27 and KP59 also had AcrAB abundances to the right of the wild-type distribution (**Fig. S1**) but in these three cases, β-lactam susceptibility was correctly predicted, based simply on β-lactamase complement (**Table 3**). KP4 is pan-β-lactam resistant due to production of CTX-M-15, NDM-1 and OXA-232 β-lactamases (**Table 3**), a phenotype that cannot be further enhanced. KP59, does not carry any acquired β-lactamase, so there is nothing to be enhanced (**Table 3**). However, AcrAB over-producing KP27 produces CTX-M-15 but has no critical errors in resistance prediction, whereas AcrAB over-producing KP13 and KP11 both produce CTX-M-15 and our predictive rules gave critical errors for cefoxitin and ertapenem (KP13), and for cefoxitin only (KP11) (**Table 3**). We suspected therefore that there must be something more than AcrAB efflux pump over-production involved in this CTX-M “enhancing” phenotype in KP13 and KP11, which was missing in KP27.

Transcription of *acrAB* is derepressed following mutations in AcrR, and activated by RamA, upon derepression of *ramA* following mutations in RamR (13, 17). WGS revealed that 6/7 AcrAB overproducing isolates (**Fig. S1, Table 3**) have a *ramR* mutation, including KP13, KP11 and the other two isolates (KP21 and KP30) where our baseline predictive rules led to critical errors for cefoxitin (**Table 3**). Importantly, however, KP27, without a CTX-M enhancing phenotype, is not a *ramR* mutant, but instead has a mutation in the AcrR binding site (17), which is the reason for AcrAB overproduction observed (**Fig. S1**).

We therefore hypothesised that the wider impact of *ramR* compared with *acrR* mutation on the proteome is responsible for the CTX-M enhancement phenotype seen in KP11 and KP13 (and also the cefoxitin critical error seen in KP21 and KP30) but not in the *acrR* mutant KP27 (**Table 3**). We aimed to identify a proteomic signature that can differentiate between *acrR* and *ramR* mutations because of this potential difference in resistance phenotype. Plotting average abundance for the YrbCDEF proteins, which are part of the RamA regulon (13, 17), across all 40 test isolates showed that the *acrR* mutant KP27 has a wild-type YrbCDEF abundance, whilst in the six *ramR* mutants, YrbCDEF abundance is higher.

To confirm our hypothesis that *ramR* and *acrR* mutation differently affect cefoxitin susceptibility, we made *acrR* or *ramR* knockout mutants of *K. pneumoniae* NCTC5055 – the strain used to establish the predictive rules (**Table S1**) – and transformed these mutants using pSU18(CTX-M-15), thereby generating an adapted set of predictive rules to be applied if *ramR* or *acrR* is lost (**Table S1**). Specifically, whether CTX-M-15 is produced or not, NCTC5055 *ramR* is cefoxitin non-susceptible, but NCTC5055 *acrR* is not. Applying these revised predictive rules corrected the critical errors for cefoxitin against the *ramR* mutants KP11, KP13, KP21, KP30, and confirmed that the *acrR* mutation in KP27 does not enhance the CTX-M phenotype (**Table 3, Table S1**). To confirm the impact of *acrR* and *ramR* mutations in *K. pneumoniae* are different, we used wild-type strain Ecl8 as background for proteomics analysis. The impact on AcrAB-TolC production was not significantly different in *acrR* or *ramR* mutatants compared with Ecl8 (t-test p>0.05, n=3), as expected from our population analysis (**Fig. S1**) but the *ramR* knockout had a wider effect on the proteome, for example, reducing OmpK35 porin production compared with the *acrR* mutant (t-test p=0.002, n=3).

### Characterisation of OmpK36 insertions in reducing carbapenem susceptibility in K. peumoniae

Factoring in *ramR* mutation removed all critical errors for cefoxitin, but a critical error remained for ertapenem in isolate KP13, suggesting that an additional mechanism is at play. Ertapenem susceptibility in CTX-M-producing *K. pneumoniae* is known to be affected by OmpK36 porin function (18-21). KP13 does not have unusually low levels of OmpK36 abundance, and there is no loss of function mutation (**Table 3**). However, OmpK36 in KP13 has a glycine-aspartate insertion after position 114 of the mature protein, which has been associated with reduced carbapenem susceptibility (22, 23). We were unable to attempt complementation of *ompK36*(GD) in KP13 due to a lack of available markers, so we constructed a series of recombinants of wild-type strain Ecl8 having, like KP13, a *ramR* mutation and carrying *bla*_CTX-M-15_ on vector pUBYT. We then disrupted *ompK36* in this recombinant and found that the derivative was ertapenem resistant, based on MIC analysis, and had reduced meropenem susceptibility, though imipenem MICs remained unchanged (**Table 4**). This is the same phenotype as expressed by clinical isolate KP13 (**Table 3**). We then complemented Ecl8Δ*ramR ompK36* pUBYT(CTX-M-15) with wild-type *ompK36* (from Ecl8) or the *ompK36*(GD) variant (generated by site directed mutagenesis from the Ecl8 wild-type allele). Ertapenem MIC was reduced 8-fold (into the susceptible range) in a transformant carrying wild-type *ompK36*, but only 2-fold (remaining non-susceptible) when carrying *ompK36*(GD). The same fold changes in meropenem MIC were also seen (**Table 4**). This confirmed that OmpK36(GD) contributes to ertapenem (but not meropenem) non-susceptibility, in a *ramR* mutant producing CTX-M-15, as seen in KP13 (**Table 3**). Accordingly, factoring in the OmpK36(GD) insertion further improves our predictive rules, and there were no remaining critical errors.

**Table 4.** MICs of carbapenems and cefepime versus *K. pneumoniae* OmpK36 variants.

| Strains | MEM | ETP | IPM | FEP |
| --- | --- | --- | --- | --- |
| Ecl8 pUBYT(CTX-M-15) | 0.125 | 0.125 | 0.5 | - |
| Ecl8$\Delta ramR$ pUBYT(CTX-M-15) | 0.125 | 1 | 0.25 | - |
| Ecl8$\Delta ramR$ *ompK36* pUBYT(CTX-M-15) pSU18 | 1 | 4 | 0.5 | - |
| Ecl8$\Delta ramR$ *ompK36* pUBYT(CTX-M-15) pSU18(*ompK36*<sub>WT</sub>) | 0.125 | 0.5 | 0.25 | - |
| Ecl8$\Delta ramR$ *ompK36* pUBYT(CTX-M-15) pSU18(*ompK36*<sub>GD</sub>) | 0.5 | 2 | 0.25 | - |
| Ecl8$\Delta ramR$ *ompK36* pUBYT(CTX-M-15) pSU18(*ompK36*<sub>DT</sub>) | 0.5 | 16 | 0.25 | - |
| Ecl8 pSU18 | - | - | - | 0.0156 |
| Ecl8 *ompK36* pSU18 | - | - | - | 0.125 |
| Ecl8 *ompK36* pSU18(*ompK36*<sub>WT</sub>) | - | - | - | 0.0312 |
| Ecl8 *ompK36* pSU18(*ompK36*<sub>GD</sub>) | - | - | - | 0.25 |
| Ecl8 *ompK36* pSU18(*ompK36*<sub>DT</sub>) | - | - | - | 0.25 |

Shaded MIC values indicate non-susceptibility according to CLSI breakpoints (36). Abbreviations: FEP, cefepime; IPM, imipenem; MEM, meropenem; ETP, ertapenem.

We looked at all the *K. pneumoniae* isolates in this study and identified two others with the *ompK36*(GD) allele. Notably, both of these carried carbapenemases, masking any effect of the GD insertion on carbapenem susceptibility. Furthermore, we identified an *ompK36* allele in the OXA-232 carbapenemase producing KP11, which generated an aspartate-threonine insertion after position 115. This *ompK36*(DT) allele has also previously been associated with reduced ertapenem susceptibility (24), but we aimed to provide the first direct experimental comparison of its effect versus the GD insertion. To do this, we used site directed mutagenesis to introduce a DT insertion into *ompK36* from Ecl8, as used for complementation before. Notably, *ompK36*(DT) was entirely unable to reduce the impact of *ompK36* disruption in Ecl8Δ*ramR* pUBYT(CTX-M-15) on ertapenem MIC, though it had the same effect as *ompK36*(GD) on meropenem MIC (**Table 4**). This suggests that OmpK36(DT) is functionally similar, and may even be more impaired, than OmpK36(GD) in the context of ertapenem susceptibility. Using Ecl8*ompK36* lacking CTX-M-15, we also showed that neither OmpK36(GD) or OmpK36(DT) could complement reduced cefepime susceptibility against Ecl8 *ompK36*. Cefepime is known to be an OmpK36 substrate (25), and this experiment confirms that the GD and DT insertions also impair cefepime entry (**Table 4**).

### Identification and characterisation of P. aeruginosa isolates that differ from the predicted pattern of β-lactam susceptibility

*P. aeruginosa* is considered very challenging for WGS-based antibiotic susceptibility prediction (9). Accordingly, we next tested a small collection of *P. aeruginosa* clinical isolates. As with *K. pneumoniae*, one of the isolates displayed a false positive for KPC using LC-MS/MS, with a low number of peptide hits. In all other cases, WGS confirmed the LC-MS/MS results. VIM β-lactamase production was seen in 3/4 isolates (**Table 5**). Analysis of the LC-MS/MS data by reference to the binary baseline predictive rules (**Table S1**) and comparison with susceptibility testing data (**Table S3**) revealed that VIM producer 301-5473 was unexpectedly aztreonam resistant. Furthermore, isolate 73-56826 did not carry any acquired β-lactamases and was therefore predicted to be susceptible to all β-lactams, but this was incorrect for ceftazidime and aztreonam (**Table 5**). According to the LC-MS/MS data, these two isolates with critical errors hyper-produce the MexAB-OprM efflux pump, relative to wild-type isolate PA01 (used to define the predictive rules). WGS revealed that this was associated with *nalC* mutation in both isolates. On the other hand, isolates 81-11963 and 404-00, carry VIM yet are aztreonam susceptible, as predicted, and do not hyper-produce MexAB-OprM, or have mutations in known regulators of its production (**Table 5**). Overproduction of MexAB-OprM in *P. aeruginosa* has been associated with aztreonam non-susceptibility in several studies of clinical isolates (26, 27). We confirmed this experimentally by knocking out *mexR* or *nalC* in *P. aeruginosa* PA01, and in both cases, the mutant was aztreonam non-susceptible (**Table S3**). Proteomics confirmed that MexA, MexB and OprM were over-produced more (1.43-fold, p=0.006; 1.58-fold, p=0.003; 1.62-fold, p=0.002, respectively for each protein) in the *mexR* mutant compared with the *nalC* mutant (**Tables S4, S5**). This has been previously shown at the level of transcription (28). None of the other 72 proteins significantly up- or downregulated upon disruption of both *mexR* and *nalC* had this same pattern of increased fold change in the *mexR* mutant (**Tables S4, S5**). We therefore conclude that the other changes in common are related second-order effects and the only direct regulatory effect common to both mutants is upregulation of the MexAB-OprM efflux pump. We were further able to differentiate these two mutations, because the ArmR protein, whose production is controlled by NalC at the level of transcription, and which is responsible for sequestering MexR and so derepressing *mexAB-oprM* transcription (28), was detectable in the proteome of the *nalC* mutant derivative PA01, but not that of the *mexR* mutant (**Tables S4, S5**).

**Table 5.** Prediction of β-lactam susceptibility in *P. aeruginosa* clinical isolates

| *P. aeruginosa*<br>Isolate ID<br>(genotype) | β-lactamases (Abundance relative to RP) | | | | | | | | | Others (Porins, efflux, AmpC) | | | | | Antibiotic Resistance Prediction | |
| --- | --- | --- | --- | --- | --- | --- | --- | --- | --- | --- | --- | --- | --- | --- | --- | --- |
|  | TEM | SHV | KPC | VIM | CMY | CTX-M | IMP | NDM | OXA-2 | OprD | MexA | MexB | OprM | AmpC | Non-susceptible | Susceptible |
| PA01 (WT) |  |  |  |  |  |  |  |  |  | 0.13 | 0.19 | 0.05 | 0.09 |  |  | ALL |
| 81-11963<br>(*oprD*) |  |  |  | 0.46 |  |  |  |  |  |  | 0.16 | 0.06 | 0.09 | 0.07 | CAZ, FEP, IPM, MEM | ATM |
| 404-00 (WT) |  |  |  | 0.71 |  |  |  |  |  | 0.19 | 0.15 | 0.05 | 0.11 |  | CAZ, FEP, IPM, MEM | ATM |
| 301-5473<br>(*nalC*, *oprD*) |  |  |  | 0.22 |  |  |  |  | 0.27 |  | 0.61 | 0.15 | 0.33 | 0.06 | CAZ, FEP, IPM, MEM | ATM |
| 73-56826 (*mpl*,<br>*nalC*) |  |  | 0.02 |  |  |  |  |  |  | 0.17 | 1.02 | 0.34 | 0.45 | 10.67 |  | CAZ, FEP, ATM,<br>IPM, MEM |

Values reported are abundance of the protein relative to the average ribosomal protein in one preparation of total cell protein. Pink highlighting denotes instances where WGS showed positive for the gene. Black/red text denotes antibiotic susceptibility phenotype that was correctly/incorrectly predicted from LC-MS/MS results.

Other protein abundance changes associated with β-lactam resistance seen in the clinical isolates included hyper-production of the chromosomal AmpC β-lactamase and reduced production of the OprD porin (27, 29). One or both of these events was seen in 3/4 test isolates within the proteomics data, but the only event consistently associated with aztreonam non-susceptibility was MexAB-OprM hyperproduction (**Table 5**). To confirm that AmpC hyper-production was not responsible; we knocked out *mpl*, encoding a UDP-muramic acid/peptide ligase whose product represses AmpC production (30), in PA01 and the mutant remained susceptible to ceftazidime, cefepime and aztreonam (**Table S3**). Remarkably, in the clinical *mpl* mutant isolate 73-65826, this hyper-produced AmpC was the second most abundant protein in the cell with an abundance 10-fold higher than the average ribosomal protein (**Table S6, Table 5**). Overall, therefore, we can adapt our β-lactam resistance predictive rules (**Table S1**) to take into consideration these important chromosomal mutations, which can be functionally confirmed through proteomics.

### Identifying isolates with atypical β-lactam susceptibility profiles directly from culture positive blood samples

Finally, we set out to find if it was possible to rapidly screen for clinical isolates that differed from the predicted β-lactam susceptibility profiles using protein extracts obtained from blood cultures via the Bruker Sepsityper kit. Among the 12 Sepsityper extracts tested using LC-MS/MS, eight were positive for *E. coli* and four were positive for *K. pneumoniae* based on ribosomal protein identification. For 8/8 of the *E. coli* and 3/4 of the *K. pneumoniae* Sepsityper extracts, we were able to correctly predict β-lactam susceptibility (**Table 6**) using binary β-lactamase identification and with reference to our predictive rules (**Table S1**). All proteomics based β-lactamase identifications matched subsequent WGS of cultured isolates (**Table 6**).

**Table 6.** Prediction of β-lactam susceptibility in Bruker Sepsityper samples.

| *E. coli*<br>Sepsityper<br>ID | 10 μg Protein Load | | | | | | | | | | |
| --- | --- | --- | --- | --- | --- | --- | --- | --- | --- | --- | --- |
| | β-lactamases | | | | Others (Porins, Efflux) | | | | Antibiotic Resistance Prediction | | |
|  | TEM | SHV | CTX-M | OXA-1 |  | OmpF | OmpC | AcrA | TolC | Non-susceptible | Susceptible |
| 69098 | 0.33 |  |  |  |  | 0.09 | 1.35 | 0.15 |  | AMOX | CXM, CRO, CTX, CAZ, FEP, ATM<br>FOX, MEM, ETP |
| 69503 |  |  |  |  |  | 0.11 | 0.54 | 0.10 | 0.09 |  | ALL |
| 69367 |  |  |  |  |  | 0.13 | 0.26 |  |  |  | ALL |
| 69357 |  |  | 1.21 | 0.07 |  |  | 1.95 | 0.11 | 0.12 | AMOX, CXM, CRO, CTX, CAZ, FEP, ATM | FOX, MEM, ETP |
| 69369 |  |  |  |  |  | 0.34 | 1.37 | 0.22 | 0.17 |  | ALL |
| 5 μg Protein Load | | | | | | | | | | | |
| 69724 |  |  | 0.84 | 0.29 |  |  | 0.38 | 0.15 | 0.21 | AMOX, CXM, CRO, CTX, CAZ, FEP, ATM | FOX, MEM, ETP |
| 69488 |  |  |  |  |  | 1.28 |  | 0.04 | 0.13 |  | ALL |
| 2.5 μg Protein Load | | | | | | | | | | | |
| 69491 | 0.12 |  |  |  |  |  | 0.29 | 0.06 | 0.16 | AMOX | CXM, CRO, CTX, CAZ, FEP, ATM<br>FOX, MEM, ETP |

| *K. pneumoniae*<br>Sepsityper<br>ID | 5 μg Protein Load | | | | | | | | | | |
| --- | --- | --- | --- | --- | --- | --- | --- | --- | --- | --- | --- |
| | β-lactamases | | | | Others (Porins, Efflux) | | | | Antibiotic Resistance Prediction | | |
|  | TEM | SHV | CTX-M | OXA-1 |  | OmpK35 | OmpK36 | AcrA | TolC | Non-susceptible | Susceptible |
| 69489 | 2.97 | 2.39 |  |  |  |  |  | 0.01 | 0.24 | AMOX | CXM, CRO, CTX, CAZ, FEP, ATM<br>FOX, MEM, ETP |
| 69694 |  |  |  |  |  |  | 0.63 | 0.02 | 0.05 | AMOX | CXM, CTX, CAZ, CRO, MEM, ET |
| 2.5 μg Protein Load | | | | | | | | | | | |
| 69815 |  |  |  |  |  |  | 1.57 | 0.09 | 0.26 | AMOX | CXM, CTX, CAZ, FEP, ATM, FO<br>CRO, MEM, ETP |
| 69723 |  |  |  |  |  |  | 1.94 | 0.04 | 0.42 | AMOX | CXM, CTX, CAZ, CRO, MEM, ET |

Values reported are abundance of the protein relative to the average ribosomal protein in one aliquot of Sepsityper extract. Pink highlighting denotes instances where WGS showed positive for the gene. Black/red text denotes antibiotic susceptibility phenotype that was correctly/incorrectly predicted from LC-MS/MS results.

*K. pneumoniae* isolate 69489 was identified via LC-MS/MS of its Sepsityper extract to express TEM-1 and SHV-1, so was predicted to only be amoxicillin non-susceptible (**Table 6**). However, susceptibility testing indicated cefoxitin and cefuroxime non-susceptibility (**Table S7, Table 6**). The LC-MS/MS data suggested TEM-1 hyperproduction and OmpK36(OmpC) porin downregulation in isolate 69489 by comparison with the two TEM-1 producing *E. coli* Sepsityper isolates, e.g. isolate 69491 (**Table 6**). These differences were confirmed following culture based proteomics for these two purified isolates, so are not an artefact of the Sepsityper extraction method (**Table S8**). WGS showed that a 9 nt deletion from the end of the Pa promoter -10 sequence of *bla*_TEM-1_ led to a strengthened Pa (8/12 identities with the perfect consensus compared with 6/12 for wild-type) and Pb (9/12 identities with the perfect consensus compared with 7/12 for wild-type), whilst retaining a wild-type P3 (**Figure S2**). This explains why TEM-1 is hyper-produced. In terms of explaining the observed OmpK36 porin downregulation in *K. pneumoniae* isolate 69489 (**Table 4, Table S8**), we noted an 885 bp region, identified as IS*1* disrupting the *ompK36* promoter.

This led to the hypothesis that disruption of *ompK36* in *K. pneumoniae* alongside over-production of TEM-1 is sufficient to confer cefoxitin and cefuroxime non-susceptibility. To test this, we knocked out *ompK36* in *K. pneumoniae* clinical isolate KP46, which according to our earlier LC-MS/MS analysis appeared to hyper-produce TEM-1 but does not have any permeability defects or carry any other β-lactamases (apart from the intrinsic SHV-1) (**Table 3**). MIC analysis of resultant recombinants (**Table S9**) supported our hypothesis.

## Conclusions

We have shown that our shotgun LC-MS/MS proteomics methodology is robust enough to facilitate prediction of antibiotic susceptibility/resistance from cultured bacteria using a single run and with reference to baseline predictive rules derived using transformants of wild-type isolates. This can even be done directly using extracts of culture positive blood. We have provided evidence that our approach can help identify isolates that sit outside of wild-type susceptibility profile, as based on our original predictive rules, focusing our attention on interesting isolates and particularly the interplay between permeability phenotypes and β-lactamase production. Finally, we have shown how this can iteratively expand our predictive rules, successfully predicting ABR more frequently and focussing down on isolates with unusual properties.

Whilst we have applied our methodology to predict β-lactam susceptibility and so study the biology of β-lactam resistance in this report, the approach would be the same for other key antibiotic classes, particularly aminoglycosides, where enzyme mediated drug modification is key to resistance in most cases. Our method has limitation when ABR involves some component of target site modification. For example in the DNA gyrase and topoisomerase IV targets of fluoroquinolones, which are subject to mutation as the major route to development of resistance to this class (14).

Whilst proteomics is limited in its ability to detect mutations, WGS is not, and there has been a lot of interest in using WGS for prediction of antibiotic susceptibility (9). However, a revolution in our ability to understand the genotype/phenotype linkage for ABR is essential if we are to use WGS direct from patient samples to correctly predict antibiotic susceptibility (9). Our proteomics methodology has the potential to dramatically improve our understanding of how ABR phenotype is influenced by genotype, bridging the gap concerning which genomic changes cause which ABR proteins to be differentially produced, and which levels of production are necessary to alter antibiotic susceptibility. Its use in parallel with WGS will therefore increase the predictive power of WGS in the future. For example, we have shown proteomic signatures that differentiate RamR and AcrR loss of function in *K. pneumoniae*, and MexR and NalC loss of function in *P. aeruginosa*. This will allow libraries to be expanded that categorise mutations – seen in WGS data – based on whether they do or do not disrupt the functions of these proteins, which will increase the future predictive power of WGS.

## Experimental

### Bacterial strains, mutants and antibiotic susceptibility testing

Transformants and mutants were made using the following strains: *K. pneumoniae* NCTC5055 (17), *K. pneumoniae* Ecl8 and Ecl8Δ*ramR* (31), *E. coli* 17 (a urinary tract isolate, and a gift from Dr. Mandy Wootton, Public Health Wales), *P. aeruginosa* PA01, and *A. baumannii* CIP 70.10. Insertional inactivation of *ramR, acrR* or *ompK36* in *K. pneumoniae* was performed using pKNOCK suicide plasmids (32). *ramR* and *acrR* DNA fragments were amplified from *K. pneumoniae* NCTC 5055 DNA by Phusion High-Fidelity DNA Polymerase (NEB, UK) with RamR_FW and RamR_KO_RV_SalI primers or AcrR_KO_FW_SalI and AcrR_KO_RV_ApaI primers. pKNOCK-Gm-*ramR* and pKNOCK-Gm-*acrR* were constructed by inserting *ramR* or *acrR* DNA fragment at SmaI/ApaI and SalI sites meanwhile pKNOCK-Gm-*ompK36* was constructed as previously described (33). *K. pneumoniae* NCTC5055 mutants were generated by electroporation whereas *ompK36* in *K. pneumoniae* Ecl8 and Ecl8Δ*ramR* were generated by conjugation. *K. pneumoniae* mutants were selected on gentamicin (5 μg.mL^-1^) and gene disruption was confirmed by PCR using RamR_R and BT87 primers for *ramR* mutants, AcrR_F and BT543 primers for *acrR* mutants and *ompK36* full-length FW and RV primers for *ompK36* mutants. Insertional inactivation of *mexR, nalC* and *mpl* in *P. aeruginosa* PA01 also used pKNOCK. DNA fragments were amplified from *P. aeruginosa* strain PA01 using primers *nalC* KO F and *nalC* KO R, *mexR* KO F and *mexR* KO R, or *mpl* KO F and *mpl* KO R. Each PCR product was then ligated into pKNOCK-GM at the SmaI site. The recombinant plasmids were transferred to *P. aeruginosa* strain PA01 by electroporation. Transformants were selected using gentamicin (30 μg.mL^-1^) and confirmed the mutations using primers *nalC* FL F and *nalC* FL R, *mexR* FL F and *mexR* FL R, or *mpl* FL F and *mpl* FL R. All primer sequences are listed in **Table S10**.

Clinical isolates were 40 human *K. pneumoniae* isolates from the collection at Severn Infection Partnership as previously described (14) and four *P. aeruginosa* (collected as part of the SENTRY antimicrobial surveillance programme and a gift from Prof Tim Walsh, Oxford University). Disc susceptibility testing was performed according to CLSI methodology (34) or microtiter broth dilution MIC testing (35) and interpreted using CLSI performance standards (36).

### Cloning β-lactamase genes, ompK36, site directed mutagenesis and transformation

All recombinant plasmids, where β-lactamase genes had been ligated into Enterobacteriales-specific vector pSU18 have already been described (13). For transformation into non-Enterobacteriales, genes were subcloned into the vector pUBYT, being the plasmid pYMAb2 (37) which we modified to remove the OXA promotor region (located upstream of the multiple cloning site) by PCR amplification using the primers pYMAb2 XbaI F and pYMAb2 XbaI R (**Table S10**), followed by digestion with XbaI and ligation to produce a circular product. Subcloning into pUBYT used the same restriction enzymes as used to originally clone the genes into pSU18 (13). Inserts were confirmed by PCR and sequencing using primers pUBYT F and pUBYT R (**Table S10**). To clone *ompK36* the gene was amplified by Phusion High-Fidelity DNA Polymerase (NEB, UK) using *K. pneumoniae* strain Ecl8 genomic DNA as template and primers OmpK36_F_SacI and OmpK36_R_HindIII (**Table S10**) The amplicon was digested with SacI and HindIII and ligated to pSU18 at the same restriction sites. Mutations encoding OmpK36(GD) and (DT) insertions were generated by two-step site-directed mutagenesis as previously (38). In brief, two PCR reactions of were set to amplify forward (M13R and GD_R primer in one, and DT_R primer in the other) and reverse (M13F and GD_F primer in one and DT_F primer in another) using pSU18(*ompK36*) as template. Each pair of PCR products was annealed and subjected to PCR using M13F and M13R in the second step. The mutant *ompK36* amplicons were digested and ligated into pSU18 at the SacI and HindIII sites. All primers are listed in **Table S10**.

Recombinant plasmids were used to transform bacteria to chloramphenicol (30 μg.mL^-1^) – for pSU18 recombinants – or kanamycin (μg.mL^-1^) – for pUBYT recombinants – resistance using electroporation as standard for laboratory-strain *E. coli*, except that for production of competent cells, *A. baumannii* cells were washed using 15% v/v glycerol in water rather than 10% v/v used for the other species.

### Preparation of samples from cultured bacteria and Sepsityper extracts

Each clinical isolate or transformant was cultured in 50 ml Nutrient Broth (Sigma) with appropriate antibiotic selection. Cultures were incubated with shaking (180 rpm) at 37°C until OD_600_ reached 0.6-0.8. Cells in cultures were pelleted by centrifugation (10 min, 4,000 × *g*, 4°C) and resuspended in 35 mL of 30 mM Tris-HCl, pH 8 and broken by sonication using a cycle of 1 sec on, 1 sec off for 3 min at amplitude of 63% using a Sonics Vibracell VC-505TM (Sonics and Materials Inc., Newton, Connecticut, USA). The sonicated samples were centrifuged at 8,000 rpm (Sorvall RC5B PLUS using an SS-34 rotor) for 15 min at 4°C to pellet intact cells and large cell debris and protein concentration in the supernatant was determined using the Bio-Rad Protein Assay Reagent according to the manufacturer’s instructions.

The processed blood culture samples (Sepsityper extracts) contained formic acid and acetonitrile. For processing on the Orbitrap LC-MS/MS, the acidity of the samples was adjusted by adding an equal volume of 5 M NaOH and three times the volume of 30 mM Tris-HCl, pH 8. Protein concentration was then estimated using the Bio-Rad Protein Dye Reagent method.

### Quantitative analysis of proteomes via Orbitrap LC-MS/MS

Total protein (varying amounts, and sometimes mixtures, as described in Results) were separated by SDS-PAGE using 11% acrylamide, 0.5% bis-acrylamide (Bio-Rad) gels and a Bio-Rad Mini-Protein Tetracell chamber model 3000×1. Gels were run at 150 V until the dye front had moved approximately 1 cm into the separating gel for ≤5 μg protein load and 4 cm for 20 μg protein load. Proteins in gels were stained with Instant Blue (Expedeon) for 5 min and de-stained in water. The 1 or 4 cm of gel lane containing each sample was cut out and proteins subjected to in-gel tryptic digestion using a DigestPro automated digestion unit (Intavis Ltd).

The resulting peptides were fractionated using an Ultimate 3000 nanoHPLC system in line with an LTQ-Orbitrap VelosPro mass spectrometer (Thermo Scientific) as we have previously described (17). The raw data files were processed and quantified using Proteome Discoverer software v1.4 (ThermoScientific) and searched against the UniProt *K. pneumoniae* strain ATCC 700721 / MGH 78578 database (5126 protein entries; UniProt accession UP000000265), the *P. aeruginosa* PAO1 database (5563 proteins; UniProt accession UP000002438), the *A. baumannii* ATCC 17978 database (3783 proteins; UniProt accession UP0006737) or the *E. coli* MG1655 database (4307 proteins; UniProt accession UP000000625), in each case, the strain-specific proteome database was augmented by addition of a mobile ABR determinant database (24694 proteins), which was generated by searching UniProt with “IncA”, “IncB” etc. to “IncZ” as the search term and downloading each list of results. Proteomic searches against the databases were performed using the SEQUEST (Ver. 28 Rev. 13) algorithm. Protein Area measurements were calculated from peptide peak areas using the “Top 3” method (39) and were then used to calculate the relative abundance of each protein. Proteins with fewer than three peptide hits were excluded from the analysis. For each sample, raw protein abundance for each protein was divided by the average abundance of the 50S and 30S ribosomal proteins also found in that sample to normalise for sample-to-sample loading variability.

### Whole genome sequencing and data analysis

Genomes were sequenced by MicrobesNG (Birmingham, UK) on a HiSeq 2500 instrument (Illumina, San Diego, CA, USA). Reads were trimmed using Trimmomatic (40) and assembled into contigs using SPAdes 3.10.1 (http://cab.spbu.ru/software/spades/). The presence of ABR genes was determined ResFinder 2.1 (41) on the Centre for Genomic Research platform (https://cge.cbs.dtu.dk/services/).

## Acknowledgements

This work was funded by grant MR/N013646/1 to M.B.A., K.J.H, O.M.W. and A.P.M. and grant MR/S004769/1 to M.B.A. from the Antimicrobial Resistance Cross Council Initiative supported by the seven United Kingdom research councils and the National Institute for Health Research. Also grant MR/T005408/1 to P.B.W. and M.B.A. from the Medical Research Council. N.S. and W.A.K.W.N.I were funded by postgraduate scholarships, respectively from the University of Bristol and the Malaysian Ministry of Education. Genome sequencing was provided by MicrobesNG.

**The authors declare no conflicts of interest**.

## Author Contributions

**Conceived the Study:** M.B.A.

**Collection of Data:** Y.T., P.D., A.S., W.A.K.W.N.I, K.J.H, supervised by A.P.M., O.M.W, P.B.W., M.B.A.

**Cleaning and Analysis of Data:** Y.T., P.D., A.S., K.J.H. supervised by M.B.A.

**Initial Drafting of Manuscript:** Y.T., M.B.A.

**Corrected and Approved Manuscript:** All Authors.

## REFERENCES

1. Didelot X, Bowden R, Wilson DJ, Peto TEA, Crook DW. 2012. Transforming clinical microbiology with bacterial genome sequencing. Nat. Rev. Genet. 13:601–12.

2. Livermore DM, Wain J. 2013. Revolutionising bacteriology to improve treatment outcomes and antibiotic stewardship. Infect. Chemother. 45:1–10.

3. Gordon NC, Price JR, Cole K, Everitt R, Morgan M, Finney J, Kearns AM, Pichon B, Young B, Wilson DJ, Llewelyn MJ, Paul J, Peto TE, Crook DW, Walker AS, Golubchik T. 2014. Prediction of Staphylococcus aureus antimicrobial resistance by whole-genome sequencing. J. Clin. Microbiol. 52:1182–91.

4. Wilson MR, Naccache SN, Samayoa E, Biagtan M, Bashir H, Yu G, Salamat SM, Somasekar S, Federman S, Miller S, Sokolic R, Garabedian E, Candotti F, Buckley RH, Reed KD, Meyer TL, Seroogy CM, Galloway R, Henderson SL, Gern JE, DeRisi JL, Chiu CY. 2014. Actionable diagnosis of neuroleptospirosis by next-generation sequencing. N. Engl. J. Med. 370:2408–17.

5. Grumaz S, Stevens P, Grumaz C, Decker SO, Weigand MA, Hofer S, Brenner T, von Haeseler A, Sohn K. 2016. Next-generation sequencing diagnostics of bacteraemia in septic patients. Genome Med. 8:73.

6. Hasman H, Saputra D, Sicheritz-Ponten T, Lund O, Svendsen CA, Frimodt-Møller N, Aarestrup FM. 2014. Rapid whole-genome sequencing for detection and characterization of microorganisms directly from clinical samples. J. Clin. Microbiol. 52:139–46.

7. Schmidt K, Mwaigwisya S, Crossman LC, Doumith M, Munroe D, Pires C, Khan AM, Woodford N, Saunders NJ, Wain J, O’Grady J, Livermore DM. 2017. Identification of bacterial pathogens and antimicrobial resistance directly from clinical urines by nanopore-based metagenomic sequencing. J. Antimicrob. Chemother. 72:104–14.

8. Stoesser N, Batty EM, Eyre DW, Morgan M, Wyllie DH, Del Ojo Elias C, Johnson JR, Walker AS, Peto TE, Crook DW. 2013. Predicting antimicrobial susceptibilities for Escherichia coli and Klebsiella pneumoniae isolates using whole genomic sequence data. J. Antimicrob. Chemother. 68:2234–44.

9. Ellington MJ, Ekelund O, Aarestrup FM, Canton R, Doumith M, Giske C, Grundman H, Hasman H, Holden MTG, Hopkins KL, Iredell J, Kahlmeter G, Köser CU, MacGowan A, Mevius D, Mulvey M, Naas T, Peto T, Rolain JM, Samuelsen Ø, Woodford N. 2017. The role of whole genome sequencing in antimicrobial susceptibility testing of bacteria: report from the EUCAST Subcommittee. Clin. Microbiol. Infect. 23:2–22.

10. Martínez-Martínez L. 2008. Extended-spectrum beta-lactamases and the permeability barrier. Clin. Microbiol. Infect. 14 Suppl 1:82–9.

11. van Boxtel R, Wattel AA, Arenas J, Goessens WHF, Tommassen J. 2016. Acquisition of Carbapenem Resistance by Plasmid-Encoded-AmpC-Expressing Escherichia coli. Antimicrob. Agents Chemother. 61:e01413–16.

12. Alekshun MN, Levy SB. 2007. Molecular mechanisms of antibacterial multidrug resistance. Cell 128:1037–50.

13. Jiménez-Castellanos JC, Wan Nur Ismah WAK, Takebayashi Y, Findlay J, Schneiders T, Heesom KJ, Avison MB. 2018. Envelope proteome changes driven by RamA overproduction in Klebsiella pneumoniae that enhance acquired β-lactam resistance. J. Antimicrob. Chemother. 73:88–94.

14. Wan Nur Ismah WAK, Takebayashi Y, Findlay J, Heesom KJ, Jiménez-Castellanos JC, Zhang J, Graham L, Bowker K, Williams OM, MacGowan AP, Avison MB. 2018. Prediction of Fluoroquinolone Susceptibility Directly from Whole-Genome Sequence Data by Using Liquid Chromatography-Tandem Mass Spectrometry To Identify Mutant Genotypes. Antimicrob. Agents Chemother. 62:e01814–17.

15. Ford PJ, Avison MB. 2004. Evolutionary mapping of the SHV beta-lactamase and evidence for two separate IS26-dependent blaSHV mobilization events from the Klebsiella pneumoniae chromosome. J. Antimicrob. Chemother. 54:69–75.

16. Foudraine DE, Dekker LJM, Strepis N, Bexkens ML, Klaassen CHW, Luider TM, Goessens WHF. 2019. Accurate Detection of the Four Most Prevalent Carbapenemases in E. coli and K. pneumoniae by High-Resolution Mass Spectrometry. Front Microbiol. 10:2760.

17. Jiménez-Castellanos JC, Wan Ahmad Kamil WN, Cheung CH, Tobin MS, Brown J, Isaac SG, Heesom KJ, Schneiders T, Avison MB. 2016. Comparative effects of overproducing the AraC-type transcriptional regulators MarA, SoxS, RarA and RamA on antimicrobial drug susceptibility in Klebsiella pneumoniae. J. Antimicrob. Chemother. 71:1820–5.

18. Mena A, Plasencia V, García L, Hidalgo O, Ayestarán JI, Alberti S, Borrell N, Pérez JL, Oliver A. 2006. Characterization of a large outbreak by CTX-M-1-producing Klebsiella pneumoniae and mechanisms leading to in vivo carbapenem resistance development. J Clin Microbiol. 44:2831–7.

19. Elliott E, Brink AJ, van Greune J, Els Z, Woodford N, Turton J, Warner M, Livermore DM. 2006. In vivo development of ertapenem resistance in a patient with pneumonia caused by Klebsiella pneumoniae with an extended-spectrum beta-lactamase. Clin Infect Dis. 42:e95–8.

20. Doumith M, Ellington MJ, Livermore DM, Woodford N. 2009. Molecular mechanisms disrupting porin expression in ertapenem-resistant Klebsiella and Enterobacter spp. clinical isolates from the UK. J Antimicrob Chemother. 63:659–67.

21. Wang XD, Cai JC, Zhou HW, Zhang R, Chen GX. 2009. Reduced susceptibility to carbapenems in Klebsiella pneumoniae clinical isolates associated with plasmid-mediated beta-lactamase production and OmpK36 porin deficiency. J Med Microbiol. 58:1196–1202.

22. Novais A, Rodrigues C, Branquinho R, Antunes P, Grosso F, Boaventura L, Ribeiro G, Peixe L. 2012. Spread of an OmpK36-modified ST15 Klebsiella pneumoniae variant during an outbreak involving multiple carbapenem-resistant Enterobacteriaceae species and clones. Eur J Clin Microbiol Infect Dis. 31:3057–63.

23. Poulou A, Voulgari E, Vrioni G, Koumaki V, Xidopoulos G, Chatzipantazi V, Markou F, Tsakris A. 2013. Outbreak caused by an ertapenem-resistant, CTX-M-15-producing Klebsiella pneumoniae sequence type 101 clone carrying an OmpK36 porin variant. J Clin Microbiol. 51:3176–82.

24. García-Fernández A, Miriagou V, Papagiannitsis CC, Giordano A, Venditti M, Mancini C, Carattoli A. 2010. An ertapenem-resistant extended-spectrum-beta-lactamase-producing Klebsiella pneumoniae clone carries a novel OmpK36 porin variant. Antimicrob Agents Chemother. 54:4178–84.

25. Tsai YK, Fung CP, Lin JC, Chen JH, Chang FY, Chen TL, Siu LK. 2011. Klebsiella pneumoniae outer membrane porins OmpK35 and OmpK36 play roles in both antimicrobial resistance and virulence. Antimicrob Agents Chemother. 55:1485–93.

26. Masuda N, Sakagawa E, Ohya S, Gotoh N, Tsujimoto H, Nishino T. 2000. Substrate Specificities of MexAB-OprM, MexCD-OprJ, and MexXY-OprM Efflux Pumps in Pseudomonas aeruginosa. Antimicrob. Agents Chemother. 44, 3322–7.

27. Castanheira M, Mills JC, Farrell DJ, Jones RN. 2014. Mutation-driven β-lactam resistance mechanisms among contemporary ceftazidime-nonsusceptible Pseudomonas aeruginosa isolates from U.S. hospitals. Antimicrob. Agents Chemother. 58, 6844–50.

28. Cao L, Srikumar R, Poole K. 2004. MexAB-OprM hyperexpression in NalC-type multidrug-resistant Pseudomonas aeruginosa: identification and characterization of the nalC gene encoding a repressor of PA3720-PA3719. Mol Microbiol. 53:1423–36.

29. Lister PD, Wolter DJ, Hanson ND. 2009. Antibacterial-resistant Pseudomonas aeruginosa: clinical impact and complex regulation of chromosomally encoded resistance mechanisms. Clin. Microbiol. Rev. 22, 582–610.

30. Calvopiña K, Avison MB. 2018. Disruption of mpl Activates β-Lactamase Production in Stenotrophomonas maltophilia and Pseudomonas aeruginosa Clinical Isolates. Antimicrob Agents Chemother. 62:e00638–18.

31. De Majumdar S, Yu J, Fookes M, McAteer SP, Llobet E, Finn S, Spence S, Monahan A, Monaghan A, Kissenpfennig A, Ingram RJ, Bengoechea J, Gally DL, Fanning S, Elborn JS, Schneiders T. 2015. Elucidation of the RamA regulon in Klebsiella pneumoniae reveals a role in LPS regulation. PLoS Pathog. 11:e1004627.

32. Alexeyev MF. (1999). The pKNOCK series of broad-host-range mobilizable suicide vectors for gene knockout and targeted DNA insertion into the chromosome of Gram-negative bacteria. Biotechniques 26:824–8.

33. Dulyayangkul P, Wan Nur Ismah Wak, Douglas EJA, Avison MB. 2020. Mutation of kvrA Causes OmpK35 and OmpK36 Porin Downregulation and Reduced Meropenem-Vaborbactam Susceptibility in KPC-Producing Klebsiella pneumoniae. Antimicrob Agents Chemother. 64:e02208–19.

34. Clinical and Laboratory Standards Institute. 2018. M02-A13. Performance standards for antimicrobial disk susceptibility tests, 13th ed. Clinical and Laboratory Standards Institute, Wayne, PA.

35. Clinical and Laboratory Standards Institute. 2015. M07-A10. Methods for dilution antimicrobial susceptibility tests for bacteria that grow aerobically; approved standard, 10th ed. Clinical and Laboratory Standards Institute, Wayne, PA.

36. Clinical and Laboratory Standards Institute. 2021. M100-S31. Performance standards for antimicrobial susceptibility testing; thirty first informational supplement. An informational supplement for global application developed through the Clinical and Laboratory Standards Institute consensus process. Clinical and Laboratory Standards Institute, Wayne, PA.

37. Kuo SC, Yang SP, Lee YT, Chuang HC, Chen CP, Chang CL, Chen TL, Lu PL, Hsueh PR, Fung CP. 2013. Dissemination of imipenem-resistant Acinetobacter baumannii with new plasmid-borne bla(OXA-72) in Taiwan. BMC Infect. Dis. 13:319.

38. Panmanee, W., Vattanaviboon, P., Eiamphungporn, W., Whangsuk, W., Sallabhan, R. and Mongkolsuk, S. 2002. OhrR, a transcription repressor that senses and responds to changes in organic peroxide levels in Xanthomonas campestris pv. phaseoli. Mol. Microbiol. 45:1647–54.

39. Silva JC, Gorenstein MV, Li GZ, Vissers JPC, Geromanos SJ. 2006. Absolute quantification of proteins by LCMSE: a virtue of parallel MS acquisition. Mol. Cell. Proteomics 5:144–156.

40. Bolger AM, Lohse M, Usadel B. 2014. Trimmomatic: a flexible trimmer for Illumina sequence data. Bioinformatics. 30:2114–20.

41. Zankari E, Hasman H, Cosentino S, Vestergaard M, Rasmussen S, Lund O, Aarestrup FM, Larsen MV. 2012. Identification of acquired antimicrobial resistance genes. J. Antimicrob. Chemother. 67:2640–4.

