## Supplementary Data File for "Identification and characterisation of *Klebsiella pneumoniae* and *Pseudomonas aeruginosa* clinical isolates with atypical β-lactam susceptibility profiles using Orbitrap liquid chromatography-tandem mass spectrometry"

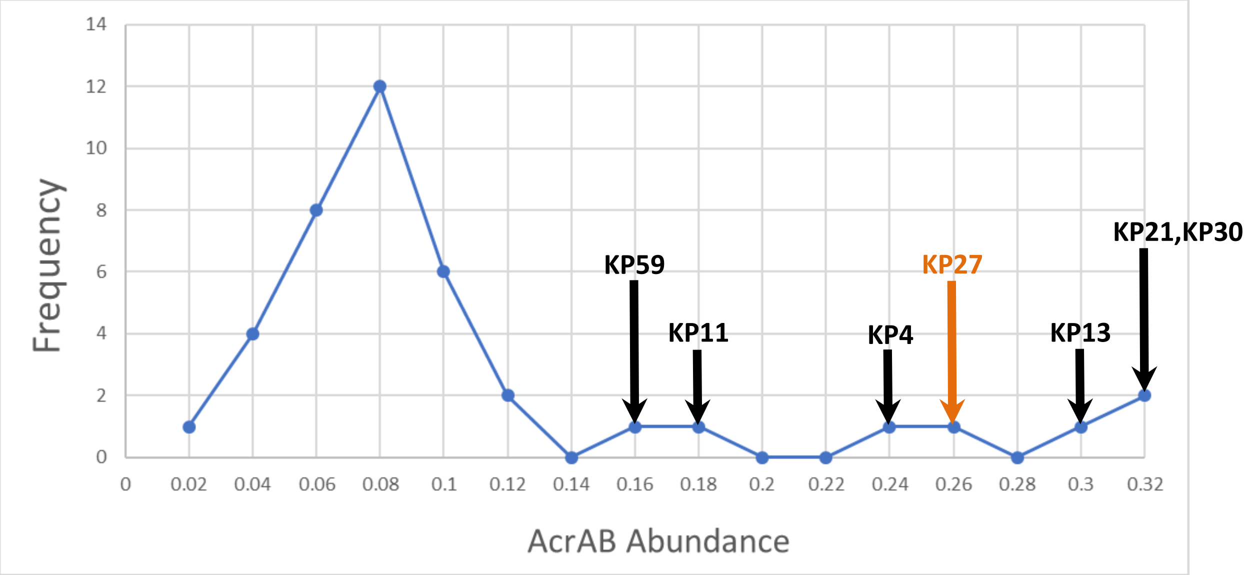

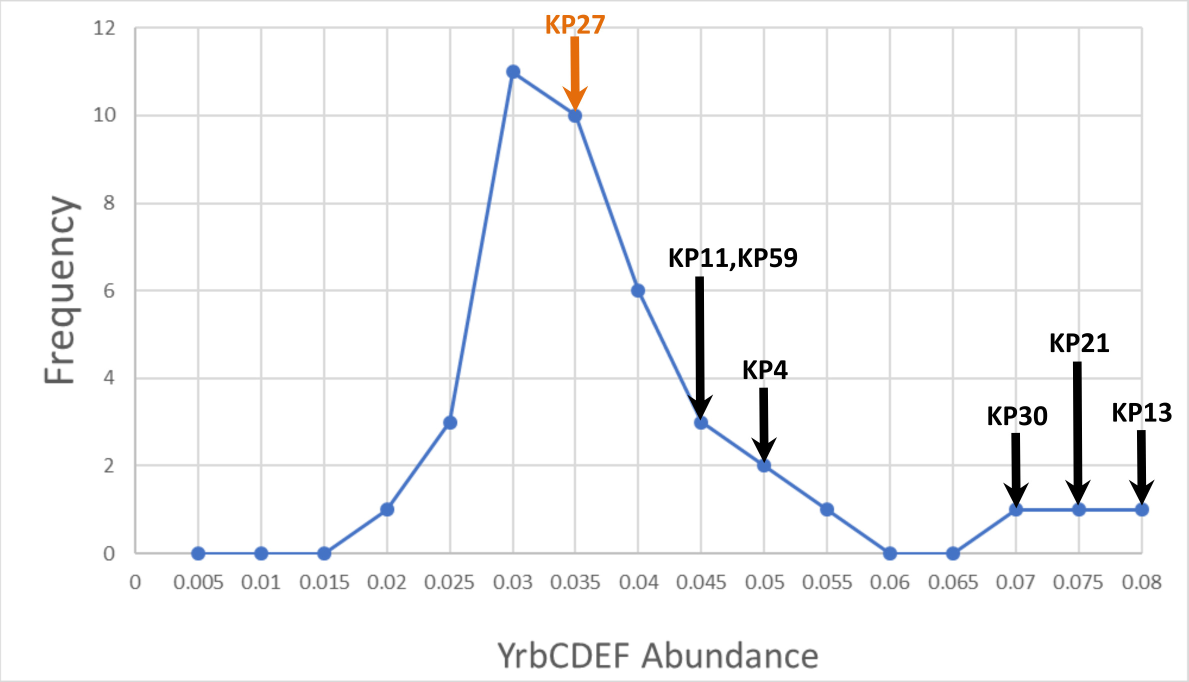

**Figure S1. Number of *K. pneumoniae* clinical isolates producing AcrAB or YrbCDEF at various abundances.**

Abundance values (normalised to ribosomal proteins) were averaged across AcrA and AcrB, or the four Yrb proteins, and the average rounded to allow bin allocation for frequency analysis.

```
                                      Pa -35                              Pb -35/Pa -10
69489      ccacggtttataaaattcttgaagacgaaagggcctcgtgatacg---------ataggt
WT         ccacggtttataaaattcttgaagacgaaagggcctcgtgatacgcctatttttataggt
           *********************************************         ******

           Pb -10
69489      taatgtcatgataataatggtttcttagacgtcaggtggcacttttcggggaaatgtgcg
WT         taatgtcatgataataatggtttcttagacgtcaggtggcacttttcggggaaatgtgcg
           ************************************************************

                                                       P3 -35                    P3 -10
69489      cggaacccctatttgtttatttttctaaatacattcaaatatgtatccgctcatgagaca
WT         cggaacccctatttgtttatttttctaaatacattcaaatatgtatccgctcatgagaca
           ************************************************************

69489      ataaccctggtaaatgcttcaataatattgaaaaaggaagagtatg
WT         ataaccctggtaaatgcttcaataatattgaaaaaggaagagtatg
           **********************************************
```

**Figure S2. Sequence of the *bla*$_{TEM-1}$ promoter in *K. pneumoniae* isolate 69489.**

Clustal Omega alignment shows identical sequence (stars) and gaps (-) caused by a 9 nt deletion in isolate 69489. The three wild-type (WT) promoters are marked: Pa/Pb in grey and P3 in green. The better consensus Pa and Pb -10 sequences generated in 69489 by the 9 nt deletion are highlighted in pink. Spacings between -35 and -10 sequence pairs are all 17 nt.

**Table S1. Antibiotic susceptibility testing for β-lactamase producing transformants.**

| Strain | FOX | CXM | CRO | CTX | CAZ | FEP | ATM | IPM | MEM | ETP |
|---|---|---|---|---|---|---|---|---|---|---|
| *K. pneumoniae* pSU18 | S | S | S | S | S | S | S | S | S | S |
| *K. pneumoniae* pSU18 KPC-3 | S | R | R | R | R | I | R | R | R | R |
| *K. pneumoniae* pSU18 VIM-1 | R | R | R | R | R | R | S | R | R | I |
| *K. pneumoniae* pSU18 IMP-1 | R | R | R | R | R | I | S | R | R | I |
| *K. pneumoniae* pSU18 OXA-48 | S | S | S | S | S | S | S | I | I | R |
| *K. pneumoniae* pSU18 CTX-M-15 | S | R | R | R | R | I | R | S | S | S |
| *K. pneumoniae* pSU18 CMY-2 | S | R | R | R | R | S | S | S | S | S |
| *K. pneumoniae* pSU18 NDM-1 | R | R | R | R | R | R | S | R | R | R |
| *K. pneumoniae acrR* pSU18 | S | S | S | S | S | S | S | S | S | S |
| *K. pneumoniae ramR* pSU18 | R | I | S | S | S | S | S | S | S | S |
| *K. pneumoniae acrR* pSU18 CTX-M-15 | S | R | R | R | R | I | R | S | S | S |
| *K. pneumoniae ramR* pSU18 CTX-M-15 | R | R | R | R | R | I | R | S | S | S |
| *E. coli* pSU18 | S | S | S | S | S | S | S | S | S | S |
| *E. coli* pSU18 KPC-3 | S | R | R | I | I | I | R | R | R | R |
| *E. coli* pSU18 VIM-1 | I | R | R | R | R | I | S | I | S | S |
| *E. coli* pSU18 IMP-1 | R | R | R | R | R | R | S | R | R | R |
| *E. coli* pSU18 OXA-48 | S | S | S | S | S | S | S | S | S | S |
| *E. coli* pSU18 CTX-M | S | R | R | R | S | I | I | S | S | S |
| *E. coli* pSU18 CMY-2 | S | R | R | R | R | S | S | S | S | S |
| *E. coli* pSU18 NDM-1 | R | R | R | R | R | R | S | R | R | R |
| *P. aeruginosa* pUBYT | | | | | S | S | S | S | S | |
| *P. aeruginosa* pUBYT KPC-3 | | | | | R | R | R | R | R | |
| *P. aeruginosa* pUBYT VIM-1 | | | | | R | R | S | R | I | |
| *P. aeruginosa* pUBYT IMP-1 | | | | | R | R | S | R | R | |
| *P. aeruginosa* pUBYT OXA-48 | | | | | S | S | S | R | I | |
| *P. aeruginosa* pUBYT CTX-M-15 | | | | | S | R | I | S | S | |
| *P. aeruginosa* pUBYT CMY-2 | | | | | S | S | S | S | S | |
| *P. aeruginosa* pUBYT NDM-1 | | | | | R | R | S | R | R | |
| *A. baumannii* pUBYT | | | S | S | S | S | | S | S | |
| *A. baumannii* pUBYT KPC-3 | | | R | R | R | R | | R | R | |
| *A. baumannii* pUBYT VIM-1 | | | R | R | R | R | | R | R | |
| *A. baumannii* pUBYT IMP-1 | | | R | R | R | R | | R | R | |
| *A. baumannii* pUBYT OXA-48 | | | I | I | S | S | | R | R | |
| *A. baumannii* pUBYT CTX-M-15 | | | R | R | I | R | | S | S | |
| *A. baumannii* pUBYT CMY-2 | | | R | R | R | S | | S | S | |

Shading: Blue, susceptible (S), Green, intermediate resistant (I), Yellow, resistant (R). Grey, no breakpoint defined by CLSI. Abbreviations: FOX, cefoxitin; CXM, cefuroxime; CRO, ceftriaxone; CTX, cefotaxime; CAZ, ceftazidime; FEP, cefepime; ATM, aztreonam; IPM, imipenem; MEM, meropenem; ETP, ertapenem.

**Table S2. Antibiotic susceptibility testing for *K. pneumoniae* clinical isolates.**

| *K. pneumoniae* Isolate ID | AST Results | | | | | | | | | |
|---|---|---|---|---|---|---|---|---|---|---|
| | FOX | CXM | CRO | CTX | CAZ | FEP | ATM | IPM | MEM | ETP |
| Kp1 | S | S | S | S | S | S | S | S | S | S |
| Kp2 | 25 | R | R | R | I | I | I | S | S | S |
| Kp3 | 24 | R | R | R | R | R | R | S | S | S |
| Kp4 | R | R | R | R | R | R | R | R | R | R |
| Kp5 | S | S | S | S | S | S | S | S | S | S |
| Kp6 | S | R | R | R | I | R | R | S | S | S |
| Kp7 | S | R | R | R | I | R | R | S | S | S |
| Kp8 | S | R | R | R | R | R | R | S | S | S |
| Kp9 | S | R | R | R | S | I | I | S | S | S |
| Kp10 | S | R | R | R | I | I | I | S | S | S |
| Kp11 | R | R | R | R | R | R | R | R | R | R |
| Kp12 | R | R | R | R | R | R | S | R | R | R |
| Kp13 | R | R | R | R | R | R | R | S | S | R |
| Kp14 | S | R | R | R | I | I | I | S | S | S |
| Kp15 | S | R | R | R | R | R | R | S | S | S |
| Kp16 | S | S | S | S | S | S | S | S | S | S |
| Kp17 | S | R | R | R | I | I | R | S | S | S |
| Kp18 | S | R | R | R | S | I | S | S | S | S |
| Kp19 | S | R | R | R | I | R | R | S | S | S |
| Kp20 | S | R | R | R | I | I | R | S | S | S |
| Kp21 | I | S | S | S | S | S | S | S | S | S |
| Kp22 | S | R | R | R | I | I | R | S | S | S |
| Kp23 | S | R | R | R | I | R | R | S | S | S |
| Kp24 | S | R | R | R | I | R | R | S | S | S |
| Kp25 | S | R | R | R | I | I | I | S | S | S |
| Kp26 | S | R | R | R | R | R | R | S | S | S |
| Kp27 | S | R | R | R | R | R | R | S | S | S |
| Kp28 | S | S | S | S | S | S | S | S | S | S |
| Kp30 | R | R | R | R | R | R | R | R | R | R |
| Kp31 | S | S | S | S | S | S | S | S | S | S |
| Kp32 | S | S | S | S | S | S | S | S | S | S |
| Kp33 | S | S | S | S | S | S | S | S | S | S |
| Kp34 | S | R | R | R | R | I | R | S | S | S |
| Kp38 | S | S | S | S | S | S | S | S | S | S |
| Kp40 | S | S | S | S | S | S | S | S | S | S |
| Kp46 | S | S | S | S | S | S | S | S | S | S |
| Kp47 | S | S | S | S | S | S | S | S | S | S |
| Kp48 | S | S | S | S | S | S | S | S | S | S |
| Kp50 | S | S | S | S | S | S | S | S | S | S |
| Kp59 | S | S | S | S | S | S | S | S | S | S |

Zone diameters are reported in mm, following standard CLSI methodology Shading: Blue, susceptible (S), Green, intermediate resistant (I), Yellow, resistant (R).

**Table S3. Antibiotic susceptibility testing for *P. aeruginosa* clinical isolates and mutant derivatives.**

| *P. aeruginosa* Isolate ID | AST Results | | | | | | | | | |
|---|---|---|---|---|---|---|---|---|---|---|
| | FOX | CXM | CRO | CTX | CAZ | FEP | ATM | IPM | MEM | ETP |
| **81-11963 (*mpl*, Δ*oprD*, VIM)** | | | | | R | R | S | R | R | |
| **404-00 (VIM)** | | | | | R | R | S | R | R | |
| **301-5473 (Δ*nalC*, Δ*oprD*, VIM)** | | | | | R | R | I | R | R | |
| **73-56826 (*mpl*, *nalC*)** | | | | | R | S | R | S | S | |
| **PA01** | | | | | S | S | S | S | S | |
| **PA01 *nalC*** | | | | | S | S | I | S | S | |
| **PA01 *mexR*** | | | | | S | S | I | S | S | |
| **PA01 *mpl*** | | | | | S | S | S | S | S | |

Shading: Blue, susceptible (S), Green, intermediate resistant (I), Yellow, resistant (R). Grey, assumed to be intrinsically resistant, so no breakpoint defined.

#### Table S4. Proteins up- and downregulated upon disruption of *mexR* in *P. aeruginosa* PA01

| Accession | Description | WT R1 | WT R2 | WT R3 | mexR R1 | mexR R2 | mexR R3 | T-test | Fold Change |
|---|---|---|---|---|---|---|---|---|---|
| E0AF19 | AaaC-A1 | | | | 1.43 | 1.68 | 1.57 | #DIV/0! | #DIV/0! |
| G3XD85 | WbpH | | | | 0.02 | 0.02 | 0.02 | #DIV/0! | #DIV/0! |
| P38108 | MucB | | | | 0.02 | 0.03 | 0.02 | #DIV/0! | #DIV/0! |
| P72158 | PurK | | | | 0.06 | 0.06 | 0.07 | #DIV/0! | #DIV/0! |
| Q00934 | PilR | | | | 0.02 | 0.02 | 0.01 | #DIV/0! | #DIV/0! |
| Q51391 | GlpR | | | | 0.01 | 0.01 | 0.02 | #DIV/0! | #DIV/0! |
| Q9HT70 | MetN | | | | 0.02 | 0.02 | 0.02 | #DIV/0! | #DIV/0! |
| Q9HTJ6 | Probable two-component response regulator | | | | 0.01 | 0.01 | 0.01 | #DIV/0! | #DIV/0! |
| Q9HUK4 | Uncharacterized protein | | | | 0.01 | 0.01 | 0.01 | #DIV/0! | #DIV/0! |
| Q9HUL8 | MutL | | | | 0.02 | 0.04 | 0.03 | #DIV/0! | #DIV/0! |
| Q9HUU1 | Oxaloacetate decarboxylase | | | | 0.03 | 0.03 | 0.04 | #DIV/0! | #DIV/0! |
| Q9HVF8 | Probable chemotaxis transducer | | | | 0.07 | 0.14 | 0.08 | #DIV/0! | #DIV/0! |
| Q9HWI2 | Probable acyl-CoA dehydrogenase | | | | 0.01 | 0.02 | 0.03 | #DIV/0! | #DIV/0! |
| Q9HY50 | Zinc-type alcohol dehydrogenase-like protein | | | | 0.02 | 0.02 | 0.03 | #DIV/0! | #DIV/0! |
| Q9HYZ3 | Probable M18 family aminopeptidase | | | | 0.02 | 0.02 | 0.02 | #DIV/0! | #DIV/0! |
| Q9HZC6 | Glycine betaine transmethylase | | | | 0.02 | 0.01 | 0.01 | #DIV/0! | #DIV/0! |
| Q9I045 | Probable two-component response regulator | | | | 0.03 | 0.01 | 0.02 | #DIV/0! | #DIV/0! |
| Q9I0V4 | Uncharacterized protein | | | | 0.68 | 0.01 | 0.59 | #DIV/0! | #DIV/0! |
| Q9I1M0 | BkdB | | | | 0.02 | 0.04 | 0.03 | #DIV/0! | #DIV/0! |
| Q9I275 | Probable glutamine synthetase | | | | 0.06 | 0.08 | 0.06 | #DIV/0! | #DIV/0! |
| Q9I2T4 | Probable binding protein component of ABC transporter | | | | 0.01 | 0.01 | 0.08 | #DIV/0! | #DIV/0! |
| Q9I4P9 | Flagellar hook protein FlgE | | | | 0.02 | 0.02 | 0.03 | #DIV/0! | #DIV/0! |
| Q9I702 | Putative 3-oxopropanoate dehydrogenase | | | | 0.02 | 0.03 | 0.03 | #DIV/0! | #DIV/0! |
| Q9I7B0 | TrkA | | | | 0.02 | 0.02 | 0.02 | #DIV/0! | #DIV/0! |
| Q51426 | RuvB | | | | 0.01 | 0.01 | 0.01 | #DIV/0! | #DIV/0! |
| Q9HXE4 | DauB | | | | 0.01 | 0.01 | 0.01 | #DIV/0! | #DIV/0! |
| Q9I371 | Probable aminotransferase | | | | 0.01 | 0.02 | 0.01 | #DIV/0! | #DIV/0! |
| Q9HUB1 | Probable chemotaxis transducer | | | | 0.08 | 0.10 | 0.08 | #DIV/0! | #DIV/0! |
| Q9HYF0 | MnmC | | | | 0.01 | 0.01 | 0.01 | #DIV/0! | #DIV/0! |
| Q9HVF0 | Uncharacterized protein | | | | 0.02 | 0.02 | 0.03 | #DIV/0! | #DIV/0! |
| Q9HY54 | FruR | 0.01 | | 0.01 | 0.06 | 0.07 | 0.07 | 0.000 | 9.042 |
| P52477 | MexA | 0.12 | 0.09 | 0.17 | 1.06 | 0.97 | 0.91 | 0.000 | 7.799 |
| Q51487 | OprM | 0.07 | 0.07 | 0.09 | 0.55 | 0.52 | 0.53 | 0.000 | 7.164 |
| P52002 | MexB | 0.07 | 0.07 | 0.08 | 0.46 | 0.51 | 0.45 | 0.000 | 6.333 |
| Q9HVZ7 | MurF | 0.01 | | 0.02 | 0.04 | 0.05 | 0.05 | 0.005 | 3.988 |
| Q9HZX9 | Probable chemotaxis transducer | 0.00 | 0.00 | 0.06 | 0.07 | 0.09 | 0.08 | 0.026 | 3.507 |
| Q9I067 | DAO domain-containing protein | 0.01 | 0.02 | | 0.03 | 0.03 | 0.03 | 0.001 | 3.346 |
| P07344 | TrpA | 0.02 | | 0.02 | 0.04 | 0.04 | 0.04 | 0.002 | 3.084 |
| Q9I0T1 | Probable acyl-CoA thiolase | 0.01 | | 0.04 | 0.06 | 0.05 | 0.05 | 0.040 | 3.027 |
| Q9HYP0 | Gln-synt_C domain-containing protein | | 0.01 | 0.02 | 0.02 | 0.03 | 0.02 | 0.036 | 2.869 |
| Q9HVA9 | Uncharacterized protein | 0.03 | 0.03 | 0.05 | 0.10 | 0.12 | 0.11 | 0.001 | 2.866 |
| Q9HZA3 | LeuC | 0.02 | 0.01 | | 0.02 | 0.03 | 0.03 | 0.015 | 2.833 |
| P34750 | Fimbrial assembly protein PilQ | 0.02 | 0.03 | 0.03 | 0.09 | 0.05 | 0.09 | 0.007 | 2.787 |
| Q9I5E4 | Aconitate hydratase | 0.01 | 0.01 | 0.03 | 0.04 | 0.06 | 0.06 | 0.010 | 2.684 |
| G3XCX1 | NarG | 0.01 | 0.01 | 0.01 | 0.03 | 0.03 | 0.03 | 0.000 | 2.664 |
| Q9I440 | Probable oligopeptidase | 0.01 | | 0.02 | 0.02 | 0.02 | 0.02 | 0.031 | 2.625 |
| Q9HZ48 | Probable binding protein component of ABC sugar transporter | 0.02 | 0.01 | 0.02 | 0.03 | 0.04 | 0.05 | 0.017 | 2.615 |
| Q9I0I4 | Probable chemotaxis transducer | 0.01 | 0.01 | 0.08 | 0.08 | 0.11 | 0.08 | 0.038 | 2.592 |
| Q9I0Z9 | Bac_luciferase domain-containing protein | 0.01 | | 0.01 | 0.01 | 0.02 | 0.01 | 0.047 | 2.570 |
| G3XD40 | Probable acyl-CoA thiolase | 0.01 | 0.01 | 0.04 | 0.04 | 0.05 | 0.04 | 0.024 | 2.435 |
| Q9HYE0 | Probable ATP-dependent RNA helicase | 0.01 | 0.01 | 0.02 | 0.04 | 0.04 | 0.03 | 0.011 | 2.362 |
| Q9HUK7 | Cyclic-di-GMP receptor FimW | 0.01 | 0.01 | 0.02 | 0.02 | 0.02 | 0.02 | 0.011 | 2.340 |
| P22567 | ArgA | 0.00 | 0.01 | 0.01 | 0.01 | 0.02 | 0.01 | 0.006 | 2.313 |
| P38100 | CarB | 0.11 | 0.08 | 0.19 | 0.23 | 0.37 | 0.26 | 0.017 | 2.307 |
| Q9HX41 | ThiD | 0.01 | | 0.01 | 0.02 | 0.02 | 0.02 | 0.037 | 2.297 |
| Q9I3M9 | CycH | 0.01 | 0.01 | 0.03 | 0.04 | 0.04 | 0.03 | 0.030 | 2.247 |
| Q9HZE0 | GdhB | 0.13 | 0.13 | 0.16 | 0.28 | 0.36 | 0.29 | 0.001 | 2.240 |
| Q9I5A5 | Pta | 0.03 | 0.02 | 0.05 | 0.06 | 0.08 | 0.08 | 0.009 | 2.232 |
| Q9I0I6 | Methyl-accepting chemotaxis protein | 0.02 | 0.03 | 0.09 | 0.09 | 0.12 | 0.09 | 0.042 | 2.227 |
| Q9HWL8 | Uncharacterized protein | 0.02 | 0.01 | 0.05 | 0.05 | 0.06 | 0.05 | 0.033 | 2.216 |
| P14532 | CcpA | 0.04 | 0.03 | 0.05 | 0.08 | 0.10 | 0.10 | 0.003 | 2.203 |
| Q9HT76 | Vitamin B12-dependent ribonucleotide reductase | 0.04 | 0.04 | 0.07 | 0.10 | 0.12 | 0.11 | 0.004 | 2.188 |
| Q9HV43 | DnaK | 0.47 | 0.41 | 1.34 | 1.35 | 1.66 | 1.80 | 0.029 | 2.165 |
| Q9HXH9 | Tgt | 0.02 | 0.01 | 0.05 | 0.04 | 0.07 | 0.06 | 0.042 | 2.154 |
| Q9HZA6 | FimV | 0.02 | 0.03 | 0.07 | 0.07 | 0.11 | 0.08 | 0.028 | 2.153 |
| Q9HW68 | Fumarate hydratase class I | 0.11 | 0.10 | 0.19 | 0.25 | 0.32 | 0.25 | 0.010 | 2.089 |
| Q9I6M5 | DavD | 0.09 | 0.07 | 0.16 | 0.20 | 0.23 | 0.25 | 0.009 | 2.088 |
| Q9HT06 | YidC | 0.05 | 0.06 | 0.10 | 0.12 | 0.15 | 0.16 | 0.013 | 2.087 |
| E1JGJ8 | PrfB | 0.02 | 0.02 | 0.06 | 0.07 | 0.09 | 0.06 | 0.037 | 2.084 |
| Q9HW06 | Probable pyrophosphohydrolase | 0.03 | 0.02 | 0.03 | 0.05 | 0.05 | 0.05 | 0.003 | 2.083 |
| Q9I271 | Uncharacterized protein | 0.02 | 0.02 | 0.05 | 0.05 | 0.07 | 0.06 | 0.038 | 2.066 |
| G3XCZ6 | FabD | 0.14 | 0.12 | 0.31 | 0.37 | 0.35 | 0.45 | 0.021 | 2.064 |
| Q9HWX4 | RibB | 0.04 | 0.02 | 0.08 | 0.09 | 0.11 | 0.09 | 0.032 | 2.039 |
| Q9I3F6 | Aer | 0.04 | 0.03 | 0.10 | 0.11 | 0.14 | 0.11 | 0.036 | 2.037 |
| Q9HYT6 | RapA | 0.03 | 0.03 | 0.07 | 0.09 | 0.09 | 0.09 | 0.006 | 2.033 |
| Q9I0L4 | Idh | 0.30 | 0.27 | 0.72 | 0.73 | 0.92 | 0.96 | 0.025 | 2.032 |
| Q9KGU7 | Dxs | 0.02 | 0.03 | 0.06 | 0.07 | 0.08 | 0.07 | 0.014 | 2.014 |

**Table S4 (continued).**

| Accession | Description | WT R1 | WT R2 | WT R3 | mexR R1 | mexR R2 | mexR R3 | T-test | Fold Change |
|---|---|---|---|---|---|---|---|---|---|
| Q9HWD0 | RpsL | 0.40 | 0.40 | 0.59 | 0.27 | 0.23 | 0.18 | 0.013 | 0.491 |
| Q9HXI9 | IscU | 0.08 | 0.08 | 0.07 | 0.05 | 0.04 | | 0.006 | 0.399 |
| P50597 | Rph | 0.04 | 0.04 | 0.03 | 0.03 | | 0.01 | 0.045 | 0.391 |
| Q9HVH7 | MscL | 0.03 | 0.05 | 0.04 | 0.03 | | 0.02 | 0.024 | 0.380 |
| Q9I1S2 | HcnB | 0.04 | 0.03 | 0.05 | 0.02 | 0.01 | 0.01 | 0.004 | 0.373 |
| Q9I367 | Uncharacterized protein | 0.03 | 0.03 | 0.03 | 0.01 | | 0.01 | 0.000 | 0.258 |
| G3XD11 | OprH | 0.02 | 0.02 | 0.02 | | | | #DIV/0! | 0.000 |
| G3XD17 | DcbC | 0.03 | 0.01 | 0.02 | | | | #DIV/0! | 0.000 |
| Q9HT10 | RsmG | 0.02 | 0.02 | 0.03 | | | | #DIV/0! | 0.000 |
| Q9HTM1 | RpoZ | 0.02 | 0.01 | 0.01 | | | | #DIV/0! | 0.000 |
| Q9HTV5 | GlpT | 0.01 | 0.02 | | | | | #DIV/0! | 0.000 |
| Q9HUK2 | ABC_trans_aux domain-containing protein | 0.01 | 0.01 | 0.03 | | | | #DIV/0! | 0.000 |
| Q9HV83 | Probable aminotransferase | 0.02 | 0.01 | 0.01 | | | | #DIV/0! | 0.000 |
| Q9HVH6 | Ferredoxin--NADP reductase | 0.01 | 0.01 | 0.01 | | | | #DIV/0! | 0.000 |
| Q9HWX2 | RibD | 0.01 | 0.01 | 0.02 | | | | #DIV/0! | 0.000 |
| Q9HXI5 | TrmJ | 0.02 | 0.01 | 0.02 | | | | #DIV/0! | 0.000 |
| Q9HYR9 | ClpP2 | 0.07 | 0.05 | 0.05 | | | | #DIV/0! | 0.000 |
| Q9I0U7 | Probable acyltransferase | 0.01 | 0.01 | 0.02 | | | | #DIV/0! | 0.000 |
| Q9I3A4 | Putative esterase | 0.01 | 0.01 | 0.01 | | | | #DIV/0! | 0.000 |
| Q9I4X2 | PqsB | 0.04 | 0.01 | 0.04 | | | | #DIV/0! | 0.000 |
| Q9I688 | Uncharacterized protein | 0.02 | 0.02 | 0.02 | | | | #DIV/0! | 0.000 |
| Q9I694 | Ribosomal RNA small subunit methyltransferase E | 0.01 | | 0.01 | | | | #DIV/0! | 0.000 |
| Q9I706 | Uncharacterized protein | 0.02 | 0.02 | 0.03 | | | | #DIV/0! | 0.000 |
| Q9I4X1 | PqsC | 0.04 | 0.02 | 0.05 | | | | #DIV/0! | 0.000 |
| Q9HVM1 | RpsT | 0.64 | 0.85 | 0.78 | | | | #DIV/0! | 0.000 |
| O30557 | AroQ1 | 0.03 | 0.04 | 0.03 | | | | #DIV/0! | 0.000 |
| Q9I347 | PrmB | 0.02 | 0.02 | 0.03 | | | | #DIV/0! | 0.000 |
| P20582 | PqsD | 0.02 | 0.02 | 0.04 | | | | #DIV/0! | 0.000 |
| P18275 | ArcD | 0.04 | 0.07 | 0.05 | | | | #DIV/0! | 0.000 |
| Q9HTQ4 | CycB | 0.04 | 0.03 | 0.02 | | | | #DIV/0! | 0.000 |
| P96963 | RadA | 0.01 | 0.01 | 0.01 | | | | #DIV/0! | 0.000 |
| Q9I4W4 | Glycine cleavage system transcriptional repressor | 0.04 | 0.03 | 0.04 | | | | #DIV/0! | 0.000 |
| Q9I351 | FolE2 | 0.04 | 0.03 | 0.04 | | | | #DIV/0! | 0.000 |
| G3XD12 | HcnC | 0.02 | 0.02 | 0.03 | | | | #DIV/0! | 0.000 |
| Q9I2S4 | Probable enoyl-CoA hydratase/isomerase | 0.02 | 0.02 | 0.03 | | | | #DIV/0! | 0.000 |
| Q9HWQ0 | Probable iron-sulfur protein | 0.03 | 0.03 | 0.02 | | | | #DIV/0! | 0.000 |
| Q9I2R7 | Probable short-chain dehydrogenase | 0.01 | 0.05 | 0.01 | | | | #DIV/0! | 0.000 |
| Q9HVX5 | Thioesterase domain-containing protein | 0.02 | 0.02 | 0.03 | | | | #DIV/0! | 0.000 |
| P25254 | Uncharacterized protein | 0.02 | 0.04 | 0.02 | | | | #DIV/0! | 0.000 |

Proteins significantly ($p<0.05$, $n=3$) upregulated >2-fold upon disruption of *mexR* are highlighted in green, those downregulated are highlighted in red. Uniprot Accession numbers of proteins similarly up- or downregulated upon disruption of *nalC* (Table S5) are highlighted in blue. Data presented are protein abundance normalised to the average of 30S and 50S ribosomal proteins in each sample. Data for biological replicates R1-3 are presented for wild type PA01 (WT) and the *mexR* mutant derivative. #DIV/0! means incalculable due to the protein being undetectable in one of the comparators.

### Table S5. Proteins up- and downregulated upon disruption of *nalC* in *P. aeruginosa* PA01

| Accession | Description | WT R1 | WT R2 | WT R3 | nalC R1 | nalC R2 | nalC R3 | T-test | Fold Change |
|---|---|---|---|---|---|---|---|---|---|
| E0AF19 | AaaC-A1 | | | | 0.89 | 0.96 | 1.08 | #DIV/0! | #DIV/0! |
| G3XD85 | WbpH | | | | 0.02 | 0.02 | 0.02 | #DIV/0! | #DIV/0! |
| P38108 | MucB | | | | 0.03 | 0.03 | 0.03 | #DIV/0! | #DIV/0! |
| P72158 | PurK | | | | 0.04 | 0.08 | 0.08 | #DIV/0! | #DIV/0! |
| Q00934 | PilR | | | | 0.02 | 0.02 | 0.02 | #DIV/0! | #DIV/0! |
| Q51391 | GlpR | | | | 0.02 | 0.01 | 0.01 | #DIV/0! | #DIV/0! |
| Q9HT70 | MetN | | | | 0.02 | 0.03 | 0.03 | #DIV/0! | #DIV/0! |
| Q9HTJ6 | Probable two-component response regulator | | | | 0.01 | 0.01 | 0.02 | #DIV/0! | #DIV/0! |
| Q9HUK4 | Uncharacterized protein | | | | 0.01 | 0.02 | 0.02 | #DIV/0! | #DIV/0! |
| Q9HUL8 | MutL | | | | 0.03 | 0.04 | 0.03 | #DIV/0! | #DIV/0! |
| Q9HUU1 | Oxaloacetate decarboxylase | | | | 0.03 | 0.04 | 0.04 | #DIV/0! | #DIV/0! |
| Q9HVF8 | Probable chemotaxis transducer | | | | 0.02 | 0.03 | 0.02 | #DIV/0! | #DIV/0! |
| Q9HWI2 | Probable acyl-CoA dehydrogenase | | | | 0.02 | 0.02 | 0.02 | #DIV/0! | #DIV/0! |
| Q9HY50 | Zinc-type alcohol dehydrogenase-like protein | | | | 0.02 | 0.02 | 0.03 | #DIV/0! | #DIV/0! |
| Q9HYZ3 | Probable M18 family aminopeptidase | | | | 0.02 | 0.02 | 0.02 | #DIV/0! | #DIV/0! |
| Q9HZC6 | Glycine betaine transmethylase | | | | 0.01 | 0.02 | 0.01 | #DIV/0! | #DIV/0! |
| Q9I045 | Probable two-component response regulator | | | | 0.03 | 0.03 | 0.03 | #DIV/0! | #DIV/0! |
| Q9I0V4 | Uncharacterized protein | | | | | 0.01 | 0.02 | #DIV/0! | #DIV/0! |
| Q9I1M0 | BkdB | | | | 0.02 | 0.03 | 0.03 | #DIV/0! | #DIV/0! |
| Q9I275 | Probable glutamine synthetase | | | | 0.05 | 0.08 | 0.07 | #DIV/0! | #DIV/0! |
| Q9I2T4 | Probable binding protein component of ABC transporter | | | | 0.01 | 0.01 | 0.01 | #DIV/0! | #DIV/0! |
| Q9I4P9 | Flagellar hook protein FlgE | | | | 0.02 | 0.03 | 0.02 | #DIV/0! | #DIV/0! |
| Q9I702 | Putative 3-oxopropanoate dehydrogenase | | | | 0.02 | 0.02 | 0.02 | #DIV/0! | #DIV/0! |
| Q9I7B0 | TrkA | | | | 0.02 | 0.03 | 0.03 | #DIV/0! | #DIV/0! |
| Q9HTL3 | RecG | | | | 0.01 | 0.36 | 0.01 | #DIV/0! | #DIV/0! |
| Q9HYQ1 | RecQ | | | | 0.01 | 0.01 | 0.01 | #DIV/0! | #DIV/0! |
| G3XD09 | GltD | | | | 0.01 | 0.01 | 0.01 | #DIV/0! | #DIV/0! |
| Q9HYX0 | Histidine kinase | | | | 0.01 | 0.01 | 0.01 | #DIV/0! | #DIV/0! |
| P57714 | MetXS | | | | 0.01 | 0.02 | 0.02 | #DIV/0! | #DIV/0! |
| Q9HWK1 | Probable acetolactate synthase large subunit | | | | 0.01 | 0.02 | 0.01 | #DIV/0! | #DIV/0! |
| Q9HZ51 | Probable ATP-binding component of ABC transporter | | | | 0.02 | 0.10 | 0.03 | #DIV/0! | #DIV/0! |
| Q9HWB0 | Probable chemotaxis transducer | | | | 0.02 | 0.02 | 0.02 | #DIV/0! | #DIV/0! |
| Q9I0Y9 | MexE | | | | 0.01 | 0.01 | 0.02 | #DIV/0! | #DIV/0! |
| C3T0J7 | TalA | | | | 0.03 | 0.03 | 0.02 | #DIV/0! | #DIV/0! |
| Q9I503 | Uncharacterized protein | | | | 0.01 | 0.01 | 0.01 | #DIV/0! | #DIV/0! |
| Q9I3L8 | Uncharacterized protein | | | | 0.03 | 0.02 | 0.02 | #DIV/0! | #DIV/0! |
| Q9HXS1 | PA3720 | | | | 0.07 | 0.06 | 0.07 | #DIV/0! | #DIV/0! |
| Q9HWK5 | PpiC2 | 0.03 | 0.41 | | 1.34 | 1.81 | 1.55 | 0.005 | 10.770 |
| P52477 | MexA | 0.12 | 0.09 | 0.17 | 0.58 | 0.75 | 0.71 | 0.000 | 5.437 |
| P22567 | ArgA | 0.00 | 0.01 | 0.01 | 0.03 | 0.04 | 0.03 | 0.001 | 4.592 |
| P45684 | RpoS | 0.01 | | 0.01 | 0.02 | 0.02 | 0.03 | 0.002 | 4.497 |
| Q51487 | OprM | 0.07 | 0.07 | 0.09 | 0.29 | 0.39 | 0.31 | 0.001 | 4.424 |
| P52002 | MexB | 0.07 | 0.07 | 0.08 | 0.25 | 0.35 | 0.30 | 0.001 | 4.002 |
| Q9HW45 | XenB | 0.02 | 0.01 | 0.04 | 0.08 | 0.09 | 0.11 | 0.003 | 3.750 |
| Q9HVZ7 | MurF | 0.01 | | 0.02 | 0.04 | 0.04 | 0.05 | 0.009 | 3.588 |
| Q9HZ48 | Probable binding protein component of ABC sugar transporter | 0.02 | 0.01 | 0.02 | 0.03 | 0.06 | 0.05 | 0.012 | 3.306 |
| Q9HYP0 | Gln-synt_C domain-containing protein | | 0.01 | 0.02 | 0.02 | 0.03 | 0.03 | 0.021 | 3.281 |
| Q9I2Q2 | MetH | 0.00 | 0.00 | 0.01 | 0.01 | 0.02 | 0.02 | 0.022 | 3.277 |
| Q9HTC2 | Probable coenzyme A transferase | 0.09 | 0.06 | 0.08 | 0.22 | 0.25 | 0.27 | 0.000 | 3.236 |
| Q9HXI2 | SecF | 0.05 | 0.22 | 0.07 | 0.31 | 0.43 | 0.36 | 0.009 | 3.206 |
| Q9I696 | ChpA | 0.01 | 0.01 | 0.11 | 0.11 | 0.14 | 0.15 | 0.032 | 3.092 |
| Q9I440 | Probable oligopeptidase | 0.01 | | 0.02 | 0.02 | 0.03 | 0.02 | 0.029 | 3.073 |
| Q9I6C2 | Probable zinc protease | 0.02 | 0.01 | 0.02 | 0.03 | 0.07 | 0.06 | 0.032 | 2.973 |
| Q9I0T1 | Probable acyl-CoA thiolase | 0.01 | | 0.04 | 0.05 | 0.05 | 0.05 | 0.046 | 2.905 |
| P34750 | Fimbrial assembly protein PilQ | 0.02 | 0.03 | 0.03 | 0.04 | 0.11 | 0.09 | 0.038 | 2.898 |
| Q9HX28 | Uncharacterized protein | | 0.01 | 0.01 | 0.01 | 0.02 | 0.02 | 0.014 | 2.775 |
| Q9I5E4 | Aconitate hydratase | 0.01 | 0.01 | 0.03 | 0.05 | 0.07 | 0.05 | 0.008 | 2.705 |
| Q9HW06 | Probable pyrophosphohydrolase | 0.03 | 0.02 | 0.03 | 0.06 | 0.07 | 0.07 | 0.001 | 2.704 |
| P14532 | CcpA | 0.04 | 0.03 | 0.05 | 0.10 | 0.12 | 0.12 | 0.001 | 2.681 |
| Q9I0L4 | Idh | 0.30 | 0.27 | 0.72 | 0.89 | 1.38 | 1.18 | 0.012 | 2.680 |
| Q9I558 | AcsA1 | 0.08 | 0.07 | 0.08 | 0.18 | 0.25 | 0.19 | 0.002 | 2.631 |
| P33642 | ThiO | 0.01 | 0.01 | 0.02 | 0.04 | 0.03 | 0.04 | 0.003 | 2.626 |
| Q9HYE0 | Probable ATP-dependent RNA helicase | 0.01 | 0.01 | 0.02 | 0.03 | 0.05 | 0.04 | 0.016 | 2.604 |
| Q9HWL8 | Uncharacterized protein | 0.02 | 0.01 | 0.05 | 0.05 | 0.07 | 0.07 | 0.021 | 2.539 |
| Q9ZN70 | Ppx | 0.03 | 0.02 | 0.03 | 0.05 | 0.08 | 0.06 | 0.006 | 2.423 |
| Q9HT76 | Vitamin B12-dependent ribonucleotide reductase | 0.04 | 0.04 | 0.07 | 0.10 | 0.14 | 0.12 | 0.005 | 2.422 |
| Q9I3M9 | CycH | 0.01 | 0.01 | 0.03 | 0.03 | 0.04 | 0.05 | 0.044 | 2.394 |
| G3XD40 | Probable acyl-CoA thiolase | 0.01 | 0.01 | 0.04 | 0.03 | 0.05 | 0.05 | 0.036 | 2.345 |
| Q9HVA9 | Uncharacterized protein | 0.03 | 0.03 | 0.05 | 0.06 | 0.11 | 0.11 | 0.017 | 2.343 |
| Q9HW68 | Fumarate hydratase class I | 0.11 | 0.10 | 0.19 | 0.27 | 0.32 | 0.33 | 0.004 | 2.341 |
| Q9HUK7 | Cyclic-di-GMP receptor FimW | 0.01 | 0.01 | 0.02 | 0.02 | 0.03 | 0.02 | 0.019 | 2.324 |
| Q9HXH9 | Tgt | 0.02 | 0.01 | 0.05 | 0.06 | 0.05 | 0.08 | 0.033 | 2.322 |
| Q9HV43 | DnaK | 0.47 | 0.41 | 1.34 | 1.39 | 1.97 | 1.75 | 0.025 | 2.301 |
| Q9HZM8 | Rne | 0.08 | 0.12 | 0.24 | 0.29 | 0.44 | 0.28 | 0.027 | 2.295 |
| Q9HT06 | YidC | 0.05 | 0.06 | 0.10 | 0.13 | 0.19 | 0.15 | 0.013 | 2.290 |
| Q9I271 | Uncharacterized protein | 0.02 | 0.02 | 0.05 | 0.05 | 0.08 | 0.07 | 0.027 | 2.287 |
| Q9HTJ1 | BetB | 0.05 | 0.04 | 0.06 | 0.11 | 0.12 | 0.12 | 0.001 | 2.275 |
| Q9I6A0 | Probable cystathionine gamma-lyase | 0.02 | 0.02 | 0.05 | 0.06 | 0.07 | 0.09 | 0.025 | 2.267 |
| Q9I4W2 | Uncharacterized protein | 0.03 | 0.03 | 0.10 | 0.10 | 0.12 | 0.14 | 0.034 | 2.258 |
| Q9HX41 | ThiD | 0.01 | | 0.01 | 0.02 | 0.02 | 0.02 | 0.040 | 2.247 |
| Q9KGU7 | Dxs | 0.02 | 0.03 | 0.06 | 0.07 | 0.09 | 0.08 | 0.012 | 2.239 |
| Q9I2T8 | PpiD PE=4 SV=1 - [Q9I2T8_PSEAE] | 0.05 | 0.05 | 0.12 | 0.14 | 0.19 | 0.17 | 0.017 | 2.224 |
| Q9I6E0 | IlvD | 0.04 | 0.04 | 0.07 | 0.09 | 0.13 | 0.11 | 0.011 | 2.212 |
| E1JGJ8 | PrfB | 0.02 | 0.02 | 0.06 | 0.07 | 0.07 | 0.08 | 0.018 | 2.210 |
| Q9HXS5 | Uncharacterized protein | 0.04 | 0.04 | 0.09 | 0.11 | 0.17 | 0.08 | 0.048 | 2.210 |
| Q9HWR5 | AMP nucleosidase | 0.01 | 0.01 | 0.02 | 0.02 | 0.04 | 0.03 | 0.011 | 2.203 |
| Q9I2Q8 | Uncharacterized protein | 0.01 | | 0.02 | 0.02 | 0.02 | 0.02 | 0.046 | 2.184 |
| Q59638 | AceF | 0.36 | 0.34 | 0.79 | 0.92 | 1.27 | 1.06 | 0.016 | 2.180 |
| Q9HZA6 | FimV | 0.02 | 0.03 | 0.07 | 0.07 | 0.11 | 0.08 | 0.021 | 2.174 |
| G3XDA2 | FabF1 | 0.09 | 0.08 | 0.17 | 0.20 | 0.29 | 0.27 | 0.015 | 2.158 |
| Q9HWX4 | RibB | 0.04 | 0.02 | 0.08 | 0.09 | 0.10 | 0.11 | 0.024 | 2.153 |
| Q9HUJ1 | Probable outer membrane protein | 0.02 | 0.02 | 0.04 | 0.05 | 0.06 | 0.05 | 0.012 | 2.136 |
| Q9HVN5 | ClpB | 0.11 | 0.12 | 0.20 | 0.24 | 0.35 | 0.33 | 0.012 | 2.136 |
| Q9I636 | GlcB | 0.13 | 0.13 | 0.19 | 0.27 | 0.38 | 0.32 | 0.004 | 2.132 |
| Q9HTM0 | SpoT | 0.01 | 0.00 | 0.01 | 0.01 | 0.02 | 0.02 | 0.034 | 2.112 |
| P31961 | Edd | 0.03 | 0.02 | 0.05 | 0.06 | 0.08 | 0.07 | 0.009 | 2.090 |
| Q9HUG8 | MsbA | 0.02 | 0.01 | 0.02 | 0.02 | 0.03 | 0.03 | 0.013 | 2.085 |
| Q9HZ65 | MtnA | 0.01 | 0.01 | 0.02 | 0.02 | 0.03 | 0.03 | 0.011 | 2.073 |
| Q9I0J6 | NuoG | 0.09 | 0.10 | 0.14 | 0.19 | 0.27 | 0.22 | 0.008 | 2.066 |
| Q51382 | HscA | 0.02 | 0.01 | 0.04 | 0.04 | 0.06 | 0.05 | 0.034 | 2.062 |
| Q9I7B7 | GlyQ | 0.05 | 0.05 | 0.06 | 0.10 | 0.10 | 0.15 | 0.014 | 2.052 |
| Q9HXB0 | PrfC | 0.03 | 0.03 | 0.05 | 0.06 | 0.08 | 0.08 | 0.011 | 2.043 |
| Q9HVZ2 | Uncharacterized protein | 0.03 | 0.02 | 0.05 | 0.06 | 0.07 | 0.07 | 0.013 | 2.039 |
| Q9HXE5 | RhlB | 0.04 | 0.03 | 0.06 | 0.08 | 0.10 | 0.08 | 0.005 | 2.037 |
| Q9HXJ8 | Der | 0.03 | 0.03 | 0.04 | 0.05 | 0.08 | 0.06 | 0.014 | 2.037 |
| Q9HUW8 | Uncharacterized protein | 0.03 | 0.02 | 0.05 | 0.06 | 0.08 | 0.07 | 0.012 | 2.035 |
| Q9HVQ3 | SpeC | 0.03 | 0.02 | 0.03 | 0.04 | 0.04 | 0.08 | 0.046 | 2.026 |
| Q9I5A5 | Pta | 0.03 | 0.02 | 0.05 | 0.05 | 0.08 | 0.07 | 0.016 | 2.019 |
| Q9HTN7 | Probable binding protein component of ABC dipeptide transpor | 0.01 | 0.01 | 0.02 | 0.02 | 0.03 | 0.03 | 0.022 | 2.017 |
| Q9I1I5 | Gcd | 0.02 | 0.01 | 0.04 | 0.04 | 0.06 | 0.05 | 0.047 | 2.011 |
| P38100 | CarB | 0.11 | 0.08 | 0.19 | 0.20 | 0.31 | 0.25 | 0.025 | 2.009 |
| P80357 | AstA | 0.02 | 0.02 | 0.03 | 0.04 | 0.05 | 0.06 | 0.008 | 2.007 |
| P27726 | Gap | 0.02 | 0.02 | 0.03 | 0.04 | 0.04 | 0.05 | 0.016 | 2.000 |

**Table S5 (continued).**

| Accession | Description | WT R1 | WT R2 | WT R3 | nalC R1 | nalC R2 | nalC R3 | T-test | Fold Change |
|---|---|---|---|---|---|---|---|---|---|
| Q9I1S2 | HcnB | 0.04 | 0.03 | 0.05 | | 0.03 | 0.02 | 0.072 | 0.434 |
| Q9HYR9 | ClpP2 | 0.07 | 0.05 | 0.05 | | 0.04 | 0.03 | 0.066 | 0.425 |
| Q9HXR1 | Uncharacterized protein | 0.03 | 0.03 | 0.04 | 0.02 | | 0.02 | 0.026 | 0.305 |
| G3XD11 | OprH | 0.02 | 0.02 | 0.02 | | | | #DIV/0! | 0.000 |
| G3XD17 | DcbC | 0.03 | 0.01 | 0.02 | | | | #DIV/0! | 0.000 |
| Q9HT10 | RsmG | 0.02 | 0.02 | 0.03 | | | | #DIV/0! | 0.000 |
| Q9HTM1 | RpoZ | 0.02 | 0.01 | 0.01 | | | | #DIV/0! | 0.000 |
| Q9HTV5 | GlpT | 0.01 | 0.02 | | | | | #DIV/0! | 0.000 |
| Q9HUK2 | ABC_trans_aux domain-containing protein | 0.01 | 0.01 | 0.03 | | | | #DIV/0! | 0.000 |
| Q9HV83 | Probable aminotransferase | 0.02 | 0.01 | 0.01 | | | | #DIV/0! | 0.000 |
| Q9HVH6 | Ferredoxin--NADP reductase | 0.01 | 0.01 | 0.01 | | | | #DIV/0! | 0.000 |
| Q9HWX2 | RibD | 0.01 | 0.01 | 0.02 | | | | #DIV/0! | 0.000 |
| Q9HXI5 | TrmJ | 0.02 | 0.01 | 0.02 | | | | #DIV/0! | 0.000 |
| Q9I0U7 | Probable acyltransferase | 0.01 | 0.01 | 0.02 | | | | #DIV/0! | 0.000 |
| Q9I3A4 | Putative esterase | 0.01 | 0.01 | 0.01 | | | | #DIV/0! | 0.000 |
| Q9I4X2 | PqsB | 0.04 | 0.01 | 0.04 | | | | #DIV/0! | 0.000 |
| Q9I688 | Uncharacterized protein | 0.02 | 0.02 | 0.02 | | | | #DIV/0! | 0.000 |
| Q9I694 | Ribosomal RNA small subunit methyltransferase E | 0.01 | | 0.01 | | | | #DIV/0! | 0.000 |
| Q9I706 | Uncharacterized protein | 0.02 | 0.02 | 0.03 | | | | #DIV/0! | 0.000 |
| O68283 | Eda | 0.02 | 0.01 | 0.02 | | | | #DIV/0! | 0.000 |
| Q51465 | FliM | 0.01 | 0.01 | 0.01 | | | | #DIV/0! | 0.000 |
| Q9HZ46 | Glk | 0.01 | 0.01 | 0.01 | | | | #DIV/0! | 0.000 |
| Q9HY41 | GlpK1 | 0.01 | 0.01 | 0.02 | | | | #DIV/0! | 0.000 |
| Q9HU42 | HisH1 | 0.01 | 0.01 | 0.02 | | | | #DIV/0! | 0.000 |
| Q9HXJ9 | Probable aminotransferase | 0.02 | 0.01 | 0.02 | | | | #DIV/0! | 0.000 |
| G3XD95 | Probable chemotaxis protein | 0.01 | 0.01 | 0.01 | | | | #DIV/0! | 0.000 |
| Q9HYY4 | Probable oxidoreductase O | 0.01 | 0.01 | 0.01 | | | | #DIV/0! | 0.000 |
| P46384 | PilG | 0.02 | 0.16 | 0.01 | | | | #DIV/0! | 0.000 |
| Q9HTF0 | Uncharacterized protein | 0.01 | 0.01 | 0.02 | | | | #DIV/0! | 0.000 |

Proteins significantly ($p<0.05$, n=3) upregulated >2-fold upon disruption of *nalC* are highlighted in green, those downregulated are highlighted in red. Uniprot Accession numbers of proteins similarly up- or downregulated upon disruption of *mexR* (Table S4) are highlighted in blue. PA3720, ArmR, is highlighted in yellow, as discussed in the text. Data presented are protein abundance normalised to the average of 30S and 50S ribosomal proteins in each sample. Data for biological replicates R1-3 are presented for wild type PA01 (WT) and the *nalC* mutant derivative. #DIV/0! means incalculable due to the protein being undetectable in one of the comparators.

#### Table S6. Proteins over-produced in Clinical isolate 73-56826 versus PA01

| Accession | Description | PAO1 R1 | PAO1 R2 | PAO1 R3 | 73 R1 | 73 R2 | 73 R3 | T-test | Fold Change |
|---|---|---|---|---|---|---|---|---|---|
| P07344 | TrpA | 0.02 | 0.00 | 0.02 | 0.13 | 0.17 | 0.11 | 0.00 | 10.18 |
| Q9HXU8 | LptF | 0.05 | 0.04 | 0.05 | 0.43 | 0.54 | 0.29 | 0.00 | 9.14 |
| Q51546 | PstB | 0.00 | 0.00 | 0.01 | 0.07 | 0.03 | 0.07 | 0.01 | 8.55 |
| P52477 | MexA | 0.12 | 0.09 | 0.17 | 0.85 | 0.58 | 1.51 | 0.02 | 7.81 |
| Q9HUE4 | MetY | 0.01 | 0.00 | 0.03 | 0.10 | 0.06 | 0.12 | 0.00 | 7.14 |
| Q9HVZ7 | MurF | 0.01 | 0.00 | 0.02 | 0.06 | 0.04 | 0.14 | 0.04 | 6.91 |
| Q51487 | OprM | 0.07 | 0.07 | 0.09 | 0.37 | 0.36 | 0.78 | 0.02 | 6.72 |
| Q9I587 | FumC1 | 0.02 | 0.02 | 0.02 | 0.07 | 0.07 | 0.19 | 0.03 | 6.35 |
| G3XDA8 | PstS | 0.07 | 0.06 | 0.08 | 0.48 | 0.13 | 0.56 | 0.03 | 5.79 |
| Q9I6M5 | DavD | 0.09 | 0.07 | 0.16 | 0.52 | 0.20 | 0.81 | 0.04 | 4.77 |
| G3XD40 | Probable acyl-CoA thiolase | 0.01 | 0.01 | 0.04 | 0.07 | 0.06 | 0.12 | 0.02 | 4.60 |
| Q9I3M9 | CycH | 0.01 | 0.01 | 0.03 | 0.07 | 0.04 | 0.12 | 0.03 | 4.60 |
| P07345 | TrpB | 0.04 | 0.04 | 0.05 | 0.21 | 0.08 | 0.28 | 0.03 | 4.48 |
| P14532 | CcpA | 0.04 | 0.03 | 0.05 | 0.29 | 0.11 | 0.15 | 0.03 | 4.31 |
| P14165 | GltA | 0.21 | 0.16 | 0.25 | 0.79 | 0.45 | 1.35 | 0.03 | 4.20 |
| P52002 | MexB | 0.07 | 0.07 | 0.08 | 0.35 | | 0.58 | 0.01 | 4.14 |
| Q9HTD9 | AdhA | 0.11 | 0.10 | 0.13 | 0.56 | 0.44 | 0.36 | 0.00 | 4.07 |
| Q9HZ48 | Probable binding protein component of ABC sugar transporter | 0.02 | 0.01 | 0.02 | 0.08 | 0.03 | 0.06 | 0.03 | 3.81 |
| Q9HTQ0 | DadA1 | 0.09 | 0.09 | 0.14 | 0.40 | 0.26 | 0.54 | 0.01 | 3.74 |
| Q9HW45 | XenB | 0.02 | 0.01 | 0.04 | 0.13 | 0.06 | 0.08 | 0.02 | 3.65 |
| Q9HU18 | DctP | 0.04 | 0.02 | 0.04 | 0.13 | 0.13 | 0.08 | 0.01 | 3.47 |
| P43904 | AroE | 0.02 | 0.01 | 0.02 | 0.05 | 0.04 | 0.06 | 0.00 | 3.26 |
| Q9HTD7 | AspA | 0.18 | 0.19 | 0.26 | 0.51 | 0.46 | 1.05 | 0.04 | 3.21 |
| Q9HZJ3 | FadA | 0.07 | 0.05 | 0.12 | 0.28 | 0.16 | 0.32 | 0.01 | 3.20 |
| Q9I5A7 | OmpA-like domain-containing protein | 0.08 | 0.05 | 0.11 | 0.20 | 0.28 | 0.26 | 0.00 | 3.11 |
| Q9HXY0 | Probable aminotransferase | 0.01 | 0.01 | 0.02 | 0.05 | 0.02 | 0.07 | 0.04 | 3.07 |
| P43334 | PhhA | 0.08 | 0.07 | 0.05 | 0.24 | 0.23 | 0.14 | 0.01 | 3.02 |
| P22008 | ProC | 0.03 | 0.01 | 0.05 | 0.08 | 0.11 | 0.06 | 0.01 | 3.01 |
| Q9I4I2 | NrdB | 0.18 | 0.20 | 0.38 | 0.57 | 0.58 | 1.13 | 0.03 | 2.98 |
| Q9HY81 | Alkyl hydroperoxide reductase C | 0.91 | 0.87 | 1.16 | 2.41 | 1.81 | 4.33 | 0.04 | 2.90 |
| Q59643 | HemB | 0.19 | 0.16 | 0.37 | 0.79 | 0.47 | 0.80 | 0.01 | 2.89 |
| P48247 | HemL | 0.13 | 0.09 | 0.17 | 0.39 | 0.21 | 0.52 | 0.03 | 2.88 |
| Q9I0D3 | CysK | 0.04 | 0.02 | 0.06 | 0.14 | 0.09 | 0.08 | 0.02 | 2.75 |
| Q9I693 | BioA | 0.02 | 0.02 | 0.03 | 0.07 | 0.05 | 0.10 | 0.03 | 2.72 |
| Q9HV70 | PanB2 | 0.04 | 0.00 | 0.05 | 0.10 | 0.09 | 0.05 | 0.04 | 2.69 |
| Q9HT95 | Acetyltransferase | 0.05 | 0.02 | 0.04 | 0.06 | 0.11 | 0.10 | 0.01 | 2.59 |
| Q9HWX4 | RibB | 0.04 | 0.02 | 0.08 | 0.15 | 0.07 | 0.13 | 0.04 | 2.53 |
| Q9HVX6 | TrpS | 0.09 | 0.08 | 0.13 | 0.18 | 0.20 | 0.38 | 0.04 | 2.53 |
| G3XCZ6 | FabD | 0.14 | 0.12 | 0.31 | 0.49 | 0.48 | 0.45 | 0.00 | 2.51 |
| Q9I4W3 | DapA | 0.08 | 0.04 | 0.14 | 0.23 | 0.27 | 0.15 | 0.03 | 2.48 |
| Q9HZA7 | AccD | 0.09 | 0.04 | 0.11 | 0.21 | 0.23 | 0.16 | 0.01 | 2.46 |
| Q9HUL7 | AmiB | 0.02 | 0.01 | 0.02 | 0.02 | 0.04 | 0.06 | 0.04 | 2.42 |
| Q9I6H7 | FliY | 0.04 | 0.02 | 0.03 | 0.09 | 0.09 | 0.06 | 0.01 | 2.41 |
| P43336 | PhhC | 0.07 | 0.06 | 0.08 | 0.22 | 0.12 | 0.17 | 0.01 | 2.36 |
| Q59641 | PpiA | 0.03 | 0.02 | 0.04 | 0.07 | 0.07 | 0.05 | 0.01 | 2.34 |
| Q9I348 | Isochorismatase domain-containing protein | 0.05 | 0.04 | 0.06 | 0.07 | 0.14 | 0.14 | 0.03 | 2.25 |
| Q9I5E2 | PrpB | 0.03 | 0.02 | 0.04 | 0.07 | 0.09 | 0.06 | 0.01 | 2.22 |
| Q9HYT1 | PBPb domain-containing protein | 0.10 | 0.08 | 0.11 | 0.23 | 0.26 | 0.15 | 0.01 | 2.22 |
| Q9I2V3 | Usp domain-containing protein | 0.05 | 0.04 | 0.06 | 0.15 | 0.11 | 0.07 | 0.04 | 2.21 |
| Q9HVF2 | Uncharacterized protein | 0.09 | 0.08 | 0.10 | 0.13 | 0.28 | 0.16 | 0.04 | 2.16 |
| Q9I020 | DUF815 domain-containing protein | 0.03 | 0.02 | 0.03 | 0.09 | 0.06 | 0.04 | 0.03 | 2.16 |
| Q60169 | HemC | 0.05 | 0.04 | 0.09 | 0.16 | 0.07 | 0.14 | 0.04 | 2.11 |
| Q9I6A8 | dITP/XTP pyrophosphatase | 0.01 | 0.01 | 0.02 | 0.02 | 0.04 | 0.03 | 0.02 | 2.10 |
| Q9HXE7 | Carboxylesterase | 0.04 | 0.03 | 0.05 | 0.08 | 0.10 | 0.07 | 0.01 | 2.08 |
| Q9HXV4 | Adk | 0.21 | 0.11 | 0.24 | 0.39 | 0.41 | 0.34 | 0.01 | 2.05 |
| Q4H482 | Beta-lactamase AmpC | | | | 13.67 | 8.26 | 9.07 | #DIV/0! | #DIV/0! |
| Q9I244 | FusB | | | | 2.67 | 0.00 | 4.61 | #DIV/0! | #DIV/0! |
| A0PCV9 | FliC | | | | 1.22 | 1.25 | 3.10 | #DIV/0! | #DIV/0! |
| Q9I653 | CreD | | | | 0.48 | 0.23 | 0.58 | #DIV/0! | #DIV/0! |
| Q9HWF3 | RpmD | | | | 0.27 | 0.22 | 0.00 | #DIV/0! | #DIV/0! |
| Q9HTL3 | RecG | | | | 0.21 | 0.00 | 0.03 | #DIV/0! | #DIV/0! |
| Q9I379 | Probable chemotaxis transducer | | | | 0.18 | 0.00 | 0.31 | #DIV/0! | #DIV/0! |
| Q9I777 | Uncharacterized protein | | | | 0.17 | 0.07 | 0.33 | #DIV/0! | #DIV/0! |
| Q9HWP3 | TyrS1 | | | | 0.13 | 0.04 | 0.13 | #DIV/0! | #DIV/0! |
| Q9HVF8 | Probable chemotaxis transducer | | | | 0.12 | 0.00 | 0.24 | #DIV/0! | #DIV/0! |
| Q9HYX3 | Probable TonB-dependent receptor | | | | 0.09 | 0.00 | 0.07 | #DIV/0! | #DIV/0! |
| Q9HXS1 | PA3720, ArmR | | | | 0.08 | 0.13 | 0.00 | #DIV/0! | #DIV/0! |

**Table S6 (continued).**

| Accession | Description | PAO1 R1 | PAO1 R2 | PAO1 R3 | 73 R1 | 73 R2 | 73 R3 | T-test | Fold Chang |
|---|---|---|---|---|---|---|---|---|---|
| Q9HUF1 | MsrA | | | | 0.08 | 0.14 | 0.13 | #DIV/0! | #DIV/0! |
| P26275 | AlgR | | | | 0.07 | 0.07 | 0.00 | #DIV/0! | #DIV/0! |
| Q9HV60 | BON domain-containing protein | | | | 0.06 | 0.06 | 0.07 | #DIV/0! | #DIV/0! |
| Q9HUL8 | MutL | | | | 0.06 | 0.00 | 0.04 | #DIV/0! | #DIV/0! |
| Q9HUU1 | Oxaloacetate decarboxylase | | | | 0.05 | 0.05 | 0.00 | #DIV/0! | #DIV/0! |
| Q9HXS0 | NalC | | | | 0.05 | 0.06 | 0.08 | #DIV/0! | #DIV/0! |
| Q9HXI6 | Serine O-acetyltransferase | | | | 0.05 | 0.06 | 0.00 | #DIV/0! | #DIV/0! |
| Q9I371 | Probable aminotransferase | | | | 0.05 | 0.00 | 0.03 | #DIV/0! | #DIV/0! |
| Q9HWI2 | Probable acyl-CoA dehydrogenase | | | | 0.05 | 0.00 | 0.13 | #DIV/0! | #DIV/0! |
| Q9HX85 | Uncharacterized protein | | | | 0.04 | 0.09 | 0.07 | #DIV/0! | #DIV/0! |
| Q9HY50 | Zinc-type alcohol dehydrogenase-like protein | | | | 0.04 | 0.02 | 0.00 | #DIV/0! | #DIV/0! |
| Q9HT52 | Probable short-chain dehydrogenase | | | | 0.04 | 0.04 | 0.00 | #DIV/0! | #DIV/0! |
| P38108 | MucB | | | | 0.04 | 0.03 | 0.00 | #DIV/0! | #DIV/0! |
| Q9X4G0 | HmgA | | | | 0.04 | 0.04 | 0.08 | #DIV/0! | #DIV/0! |
| Q9HW73 | Uncharacterized protein | | | | 0.04 | 0.03 | 0.00 | #DIV/0! | #DIV/0! |
| Q9I4L2 | DUF2025 domain-containing protein | | | | 0.03 | 0.06 | 0.00 | #DIV/0! | #DIV/0! |
| Q9I5X8 | Methyltransf_11 domain-containing protein | | | | 0.03 | 0.03 | 0.00 | #DIV/0! | #DIV/0! |
| Q9HUT3 | Probable bacterioferritin | | | | 0.03 | 0.08 | 0.07 | #DIV/0! | #DIV/0! |
| Q9I2A2 | FahA | | | | 0.03 | 0.00 | 0.03 | #DIV/0! | #DIV/0! |
| Q9HV90 | PhuT | | | | 0.03 | 0.01 | 0.00 | #DIV/0! | #DIV/0! |
| Q9HTY0 | Probable secretion pathway ATPase | | | | 0.03 | 0.00 | 0.11 | #DIV/0! | #DIV/0! |
| Q9HZI6 | Probable soluble lytic transglycosylase | | | | 0.03 | 0.00 | 0.03 | #DIV/0! | #DIV/0! |
| Q9HWK1 | Probable acetolactate synthase large subunit | | | | 0.02 | 0.00 | 0.06 | #DIV/0! | #DIV/0! |
| Q51391 | GlpR | | | | 0.02 | 0.03 | 0.00 | #DIV/0! | #DIV/0! |
| Q9I702 | Putative 3-oxopropanoate dehydrogenase | | | | 0.02 | 0.02 | 0.07 | #DIV/0! | #DIV/0! |
| Q9HU78 | HutC | | | | 0.02 | 0.03 | 0.00 | #DIV/0! | #DIV/0! |
| Q9HVW1 | Probable ATP-binding component of ABC transporter | | | | 0.02 | 0.02 | 0.00 | #DIV/0! | #DIV/0! |
| Q9I2T4 | Probable binding protein component of ABC transport | | | | 0.02 | 0.00 | 0.03 | #DIV/0! | #DIV/0! |
| Q9HZK2 | Uncharacterized protein | | | | 0.02 | 0.03 | 0.00 | #DIV/0! | #DIV/0! |
| Q9I4P9 | FlgE | | | | 0.02 | 0.02 | 0.05 | #DIV/0! | #DIV/0! |
| Q9HUE2 | Uncharacterized protein | | | | 0.02 | 0.02 | 0.00 | #DIV/0! | #DIV/0! |
| Q9HZ27 | MlaD domain-containing protein | | | | 0.02 | 0.02 | 0.00 | #DIV/0! | #DIV/0! |
| Q9HU13 | Nudix hydrolase domain-containing protein | | | | 0.01 | 0.02 | 0.00 | #DIV/0! | #DIV/0! |
| P72161 | PbpG | | | | 0.01 | 0.02 | 0.00 | #DIV/0! | #DIV/0! |
| Q9HUW7 | Probable two-component response regulator | | | | 0.01 | 0.00 | 0.03 | #DIV/0! | #DIV/0! |
| Q9HYX0 | Histidine kinase | | | | 0.01 | 0.00 | 0.01 | #DIV/0! | #DIV/0! |
| Q9HTJ6 | Probable two-component response regulator | | | | 0.01 | 0.01 | 0.00 | #DIV/0! | #DIV/0! |
| Q9HZ93 | 2-dehydro-3-deoxy-phosphogluconate aldolase | | | | 0.01 | 0.02 | 0.00 | #DIV/0! | #DIV/0! |
| Q9HX07 | Mpl | | | | 0.00 | 0.02 | 0.06 | #DIV/0! | #DIV/0! |
| Q9I0V4 | Uncharacterized protein | | | | 0.00 | 0.01 | 0.03 | #DIV/0! | #DIV/0! |
| Q9HX91 | Uncharacterized protein | | | | 0.00 | 0.03 | 0.05 | #DIV/0! | #DIV/0! |

Proteins significantly ($p<0.05$, $n=3$) upregulated >2-fold in clinical isolate 73-56826 (*nalC*, *mpl*) are highlighted in green, Proteins discussed in the text: MexA, MexB, OprM, AmpC and PA3720, ArmR, are highlighted in yellow. Data presented are protein abundance normalised to the average of 30S and 50S ribosomal proteins in each sample. Data for biological replicates R1-3 are presented for wild type PA01 (WT) and clinical isolate 73-56826 (73) #DIV/0! means incalculable due to the protein being undetectable in one of the comparators.

**Table S7. Antibiotic susceptibility results for Sepsityper isolates.**

| MALDI Sepsityper Isolates | AMOX | FOX | CXM | CRO | CTX | CAZ | FEP | ATM | MEM | ETP |
|---|---|---|---|---|---|---|---|---|---|---|
| ***E. coli* 69098** | R | S | S | S | S | S | S | S | S | S |
| ***E. coli* 69503** | S | S | S | S | S | S | S | S | S | S |
| ***E. coli* 69367** | S | S | S | S | S | S | S | S | S | S |
| ***E. coli* 69357** | R | S | R | R | R | R | R | R | S | S |
| ***E. coli* 69369** | S | S | S | S | S | S | S | S | S | S |
| ***E. coli* 69724** | R | S | R | R | R | R | I | I | S | S |
| ***E. coli* 69488** | S | S | S | S | S | S | S | S | S | S |
| ***E. coli* 69491** | R | S | S | S | S | S | S | S | S | S |
| ***K. pneumoniae* 69489** | R | I | I | S | S | S | S | S | S | S |
| ***K. pneumoniae* 69694** | R | S | S | S | S | S | S | S | S | S |
| ***K. pneumoniae* 69815** | R | S | S | S | S | S | S | S | S | S |
| ***K. pneumoniae* 69723** | R | S | S | S | S | S | S | S | S | S |

Shading: Blue, susceptible (S), Green, intermediate resistant (I), Yellow, resistant (R).

(Data are from the clinical diagnostic laboratory, but were confirmed using disc testing of cultured isolates)

**Table S8. Culture-based LC-MS/MS data on selected Sepsityper isolates.**

| Isolate ID | β-lactamases | | | | Porins, Efflux | | | | |
|---|---|---|---|---|---|---|---|---|---|
| | TEM | SHV | CTX-M | OXA-1 | | OmpF/K35 | OmpC/K36 | AcrA | TolC |
| ***E. coli* 69491** | 0.07 | | | | | 0.04 | 2.11 | 0.10 | 0.10 |
| ***K. pneumoniae* 69489** | 0.80 | 0.29 | | | | 0.07 | 0.05 | 0.09 | 0.05 |

Values reported are abundance of the protein relative to the average 30S and 50S ribosomal protein in one preparation of total cell protein. Pink highlighting denotes instances where WGS showed positive for the gene.

**Table S9. MICs (μg.ml$^{-1}$) against *K. pneumoniae* recombinants and mutants**

| | Cefuroxime (CXM) | Cefoxitin (FOX) |
|---|---|---|
| KP46 (TEM-1 hyper-producer) | 4 | 4 |
| KP46 *ompK36* | 32 | 64 |

The grey highlight indicates non-susceptible based on CLSI criteria.

**Table S10. Primers used in this study.**

| Primer | Sequence (5'-3') |
|---|---|
| **pYMAb2 XbaI F** | ATGACTTCTAGACAGCAAATGG |
| **pYMAb2 XbaI R** | GAGATCTCTAGATTAACCGTTC |
| **pUBYT F** | GCAAGAAGGTGATGAATCTACA |
| **pUBYT R** | GTGGCAGCAGCCAACTCA |
| **RamR_KO_FW** | GCGATGAAAGTCGTCAAGACGATT |
| **RamR_KO_RV_SalI** | AAAGTCGACGAAAGCCGCGGTGGCAGCTT |
| **RamR_R** | CGCCCCAGTCGATATAGCTGT |
| **AcrR_KO_FW_SalI** | AAAGTCGACGTGAAACCCGGCAACTGATT |
| **AcrR_KO_RV_ApaI** | AAAGGGCCCACTGAGAGTGGATCGTTGGG |
| **AcrR F** | ACCTCGAGTGTCCAATTTCAAAATGTTC |
| **OmpK36 full-length FW** | GAGGCATCCGGTTGAAATAG |
| **OmpK36 full-length RV** | ATTAATCGAGGCTCCTCTTAC |
| **OmpK36_F_SacI** | AAAGAGCTCTTAGTGCGTATTTCCCTGAC |
| **OmpK36_R_HindIII** | TATAAGCTTTTGTTATGCAGCTTGCAACTT |
| **GD_F** | GAATTCGGCGGCGACGGCGACACCTACGGTTCT |
| **GD_R** | ACCGTAGGTGTCGCCGTCGCCGCCGAATTCCGG |
| **DT_F** | TTCGGCGGCGACACCGACACCTACGGTTCTGAC |
| **DT_R** | AGAACCGTAGGTGTCGGTGTCGCCGCCGAATTC |
| **BT87** | CACTTAACGGCTGACATGG |
| **BT543** | TGACGCGTCCTCGGTAC |
| **M13F** | GTAAAACGACGGCCAGT |
| **M13R** | CAGGAAACAGCTATGAC |
| ***nalC* KO F** | GCCCACATGATCGGGGAAAT |
| ***nalC* KO R** | CCCCTGCTCGTAGAAGGACT |
| ***mexR* KO F** | AGCTTATCGACGAACAACGC |
| ***mexR* KO R** | GGGCAAACAACTCGTCATGC |
| ***mpl* KO F** | CCGCAGAACTTCGGGGTTT |
| ***mpl* KO R** | GAGGTCGGGGAAGATATCCG |
| ***nalC* FL F** | GAGCGGAAGTGCTTGCCAAA |
| ***nalC* FL R** | CTGCTCGAACGTACCCTGCC |
| ***mexR* FL F** | AGGTTTACTCGGCCAAACCAA |
| ***mexR* FL R** | ATGTTCTTAAATATCCTCAAGCGGT |
| ***mpl* FL F** | AATCTGCCGCCCATTCACAG |
| ***mpl* FL R** | TCATCGCGCCCCTCACTC |
